# Pioneer and Altimeter: Rapid and Scalable Analysis of DIA Proteomics Data Optimized for Narrow Isolation Windows

**DOI:** 10.64898/2026.02.16.706201

**Authors:** Nathan T. Wamsley, Emily M. Wilkerson, Michael B. Major, Dennis Goldfarb

## Abstract

The scale of mass spectrometry–based proteomics studies has surged due to faster instrumentation and automated sample preparation, producing terabytes of data across thousands of files and pushing analysis software to their limits. Here we introduce Pioneer and Altimeter for rapid and scalable analysis of data-independent acquisition (DIA) proteomics data. Altimeter generates in silico spectral libraries, which Pioneer uses to identify and quantify peptides and proteins. Pioneer exceeds peptide coverage by up to 10%, improves quantitative accuracy, and runs four to eight times faster than state-of-the-art tools. Using 16 CPU cores and 64 GB of memory, it searched nearly 10,000 yeast runs in 60 hours and 2,200 human runs in 24 hours. Both tools model isolation-window effects on fragment isotopes to exploit narrow windows on fast-scanning time-of-flight instruments. We validated false-discovery-rate and false-transfer-rate control across diverse instruments, sample types, and acquisition settings. Pioneer is cross-platform, with graphical and command-line interfaces.

## 1 Introduction

Mass spectrometry (MS)-based proteomics enables high-throughput quantitation of peptides and proteins from complex biological samples. Increasingly for quantitative proteomics, these data have been acquired by data-independent acquisition (DIA). Faster and more sensitive mass analyzers have grown data output by more than an order of magnitude in the past decade but without a corresponding increase in the speed of downstream analysis tools [1, 2]. Co-fragmentation of multiple precursors further complicates that analysis [3–6]. To mitigate this problem the fastest instruments acquire data with narrow isolation windows to reduce co-isolation and scan complexity, but doing so distorts the relative fragment isotope abundances [7]. Current analysis methods struggle to keep pace with proteomics data production and do not exploit the unique properties of narrow isolation windows.

Genetic screens, population-level clinical studies, single-cell experiments, and other large analyses include hundreds of samples [8–13]. The latest instruments process these samples by acquiring high-sensitivity and high-mass-accuracy scans at up to 270 Hz to produce hundreds of gigabytes in MS data files per day [14–16]. Ideally the investigators could perform these analyses at least several times faster than data acquisition and with only modest personal computers.

A further, largely ignored challenge of contemporary DIA methods is that narrow isolation windows distort fragment-ion isotope distributions relative to their unconditional expected profiles. Most DIA analysis tools rely on empirical or predicted spectral libraries [17–24]. In either case these libraries ultimately depend on previous experiments where the isolation windows were centered on different precursor features as compared to the down-stream DIA acquisition method [25–28]. This leads to discrepancies between the library spectra and the DIA data under analysis [7, 29, 30]. Existing analysis tools neither model nor correct for these discrepancies.

Here we introduce Pioneer and Altimeter, two free and open-source tools for fast analysis of DIA proteomics data. Pioneer is responsible for peptide and protein group identification and quantification, and Altimeter is a deep learning model that predicts total relative fragment intensity. Pioneer adapts predicted spectra to different isolation windows without fine-tuning, transfer learning, or re-prediction. Moreover, Pioneer improves analysis speed over state-of-the-art software by a factor of four to eight without sacrificing identifications or quantitative precision. We first present a general overview of the Pioneer pipeline and the Altimeter model. Next, we demonstrate using several benchmark datasets and extensive entrapment analyses that Pioneer and Altimeter provide a framework for DIA analysis that keeps pace with the fastest mass spectrometers.

## 2 Results

### 2.1 A unified approach to isotope-aware DIA analysis

We developed Pioneer to perform fast DIA analysis and to account for imperfect precursor isolation (Fig. 1A, Extended Data Fig. 1). This was motivated by the fact that isolating different subsets of a precursor’s isotopic envelope produces MS2 spectra with distinct intensity patterns (Fig. 1B). While a fragment’s total relative intensity does not change, the transmission of different precursor isotopes redistributes signal among the fragment’s various isotopes. This redistribution leads to deviations from the expected fragment ratios learned by models trained on DDA data. With the widespread adoption of narrow isolation windows, these effects are no longer rare edge cases: roughly a third of doubly charged precursors are substantially split between adjacent windows at a width of 2 m/z (Fig. 1C).

**Fig 1:**
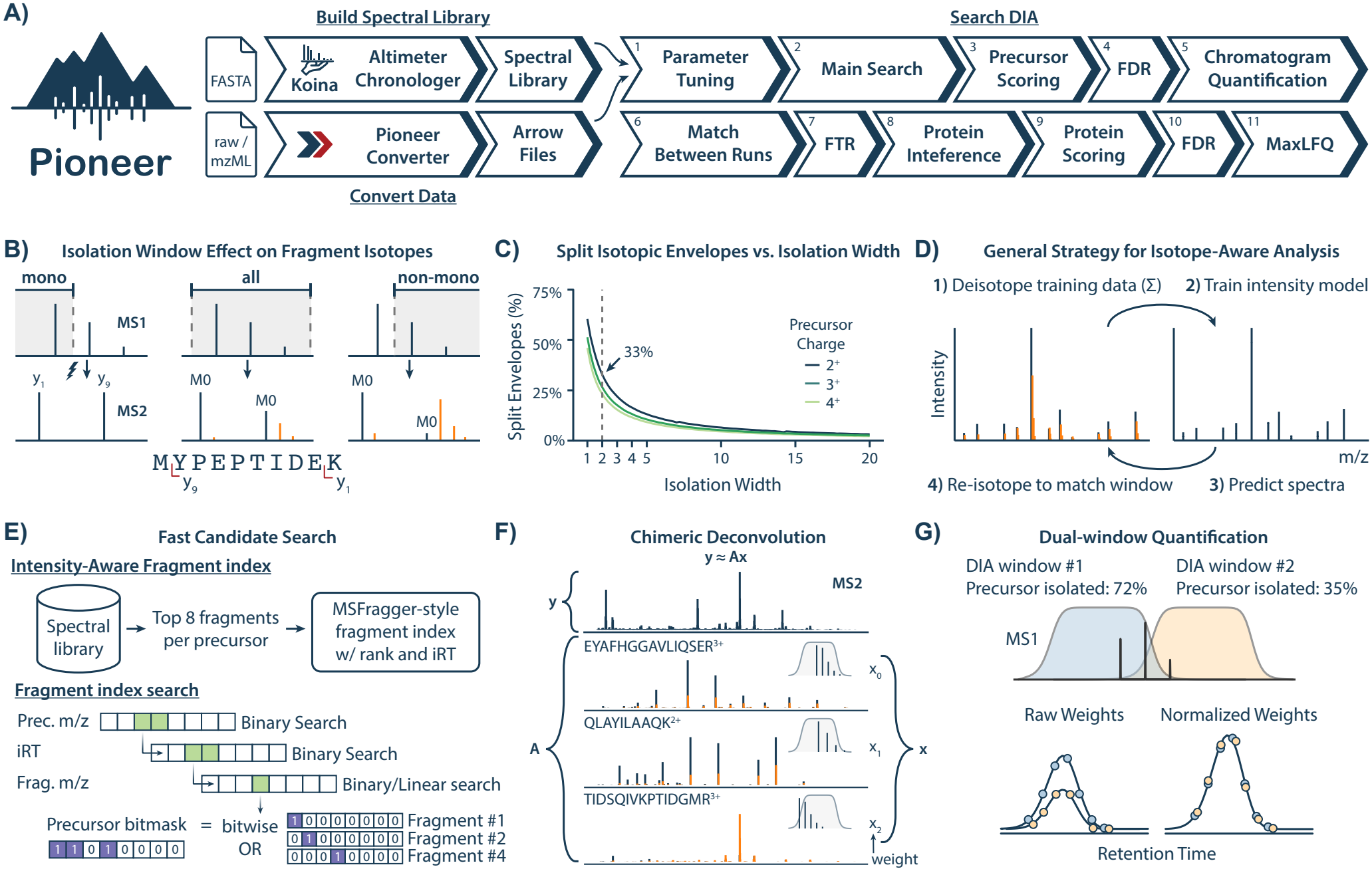
Pioneer workflow for DIA analysis. **A)** Overall pipeline for building predicted spectral libraries, converting data, and searching DIA runs. Chronologer predicts retention times [31]. MaxLFQ is used to infer protein abundances from peptide abundances [32]. False discovery (FDR) and false transfer rates (FTR) are controlled. **B)** Example of how isolation windows that capture the monoisotopic, all, or non-monoisotopic precursors alter fragment isotope distributions. Non-monoisotopic fragment peaks are shown in orange. **C)** Theoretical frequency of splitting a precursor’s isotopic envelope on a tryptic human digest where both windows isolate ≥25% of the envelope. **D)** General strategy for isotope-aware analysis. Training spectra are deisotoped (summed) and used to train Altimeter, a fragmentation model that predicts total fragment intensity. The Altimeter predictions are subsequently re-isotoped to match experimental isolation windows. **E)** An intensity- and iRT-aware fragment index is constructed during spectral library prediction using the top fragments per precursor. Candidate precursors are scored by combining 8-bit fragment bitmasks with a bitwise OR to generate a precursor bitmask. Precursor bitmask patterns enriched for targets undergo an initial classification step to distinguish targets from decoys. **F)** Observed spectra are modeled as a linear combination of predicted spectra from candidate precursors while minimizing Huber loss. Predictions are re-isotoped to the acquisition window (inset shows window position relative to the precursor). The resulting weights are used for quantification. **G)** Dual-window quantification using precursor isolation percentages to normalize fragment intensities. Quadrupole transmission profiles are fit during parameter tuning. Chromatograms are smoothed by a Whittaker–Henderson filter [33, 34].

Pioneer addresses this mismatch by using spectral libraries generated by Altimeter (deployed on the Koina server [35]), which predicts total fragment intensities rather than monoisotopic intensities. Predicted spectra can then be re-isotoped on a per-scan basis according to a fitted quadrupole transmission profile, allowing for direct adaptation to the acquisition window (Fig. 1D).

To efficiently identify candidate precursors, Pioneer uses an intensity-aware fragment index that restricts each MS2 spectrum to a set of plausible precursor matches (Fig. 1E). The index, inspired by MSFragger [36], contains only the eight most intense predicted fragments per precursor. Each fragment is assigned an 8-bit bitmask that encodes its intensity rank within the precursor’s library fragment spectrum. For a given MS2 spectrum, every precursor accumulates a bitmask, initialized to zero and updated by a bitwise OR with the bitmask of each matching fragment. A rapid calibration step identifies which of the resulting 256 bit patterns are enriched for targets; once all fragments in the scan have been queried, a precursor passes the fragment index for that scan if its accumulated vector matches one of these target-enriched patterns.

To further refine candidate precursors and perform quantification, Pioneer models each MS2 spectrum as a linear combination of re-isotoped library spectra using robust regression (Fig. 1F). This formulation accounts for chimeric fragmentation by assigning weights to co-isolated precursors. Because each scan may isolate only a portion of a precursor’s isotopic envelope, Pioneer normalizes weights by the isolated fraction to recover precursor-level intensity and place adjacent windows on a common scale. After correction, Pioneer performs dual-window quantification, combining weights from adjacent windows into a single chromatographic trace with twice the data points (Fig. 1G). Together, these components prevent changes in isolation windows from altering precursor quantification.

### 2.2 Decoupled, continuous prediction of total fragment intensity

DIA analysis benefits from predictive models that can be optimized for each dataset and, ideally, each run. However, adapting predictions to run-specific collision energies or isolation windows typically requires repeated neural network inference or fine-tuning; both are computationally expensive. We therefore designed Altimeter to decouple run-specific adaptation from model evaluation through a representation that can be adjusted after inference (Fig. 2A).

**Fig 2:**
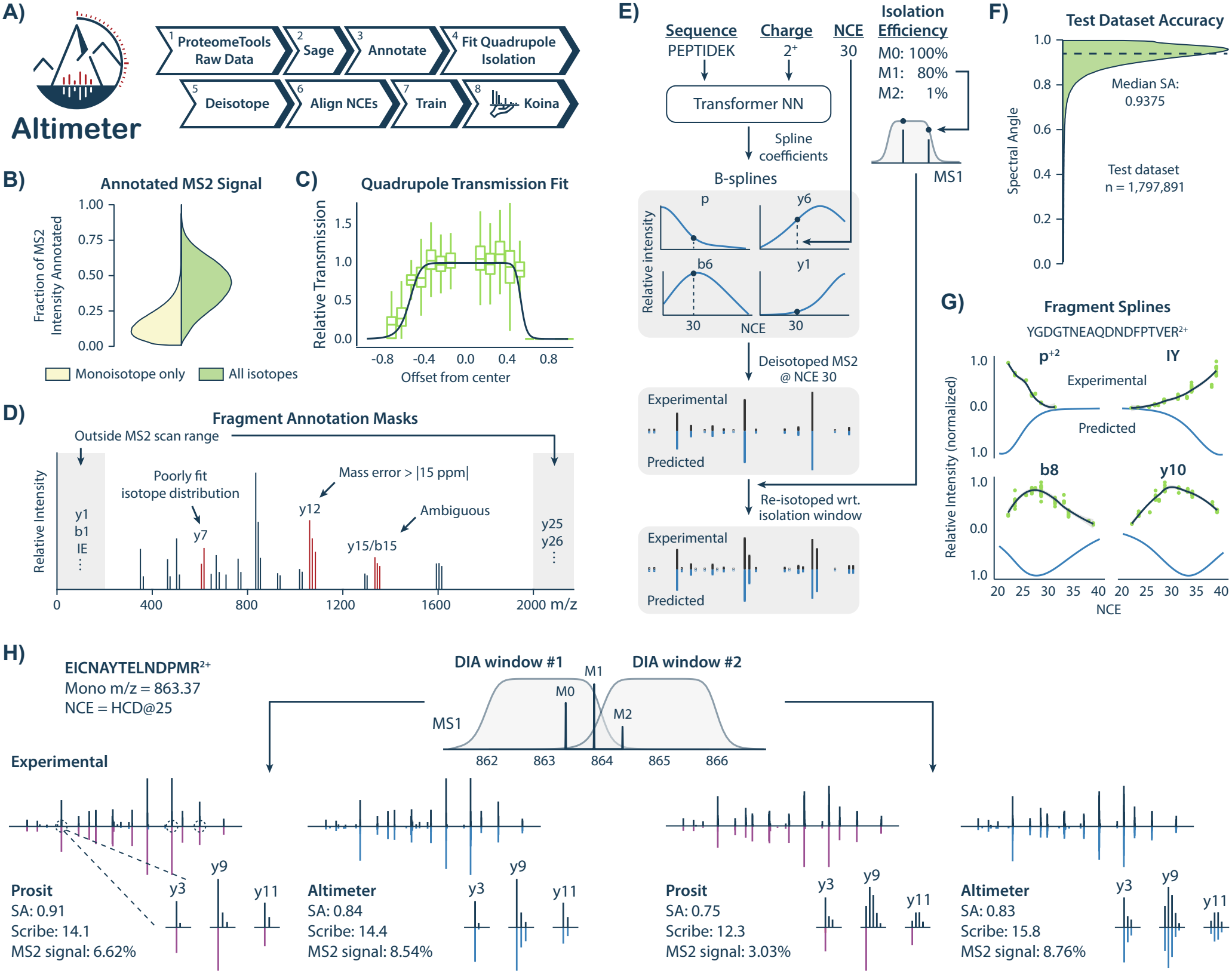
Altimeter deep learning model for peptide fragmentation prediction. **A)** Model development workflow, from raw spectra to training and deployment. **B)** Fraction of MS2 spectrum intensity annotated in the training data, with and without higher isotopes. **C)** Quadrupole transmission profile fit for a set of runs having the same quadrupole calibration date. Transmission is relative to the central isotope in the window. Boxplots represent estimated transmission rates for neighboring isotopes, calculated from their relative MS2 contribution compared to theoretical precursor isotope abundances. **D)** Cartoon schematic of fragment annotation masks: outside scan range, isotope misfit, high mass error, and ambiguous assignments. **E)** Schematic of the Altimeter model: peptide sequence and charge are input to a transformer network, which outputs B-spline coefficients that model intensity profiles across normalized collision energies (NCE). Monoisotopic spectra are predicted by evaluating the splines at the desired NCE. Spectra can be further re-isotoped given the relative isolation efficiency of each precursor isotope. **F)** Spectral angle accuracy distribution on the held-out test dataset. **G)** Example fragment B-splines for a peptide from the test set, showing experimental (above) and Altimeter-predicted (below). Intensities and spline predictions were rescaled by the maximum across NCEs for visualization. **H)** Example prediction for a peptide precursor spanning two 2 m/z-wide isolation windows on an Orbitrap Astral. Experimental and predicted spectra are compared for Altimeter and Prosit, with spectral angle (SA), Scribe score, and percentage of annotated MS2 signal indicated.

As an initial step, we reprocessed ProteomeTools [37, 38] to account for precursor isolation and fragment isotopes. The addition of higher isotopes doubled the annotated MS2 signal compared to monoisotopic peaks (Fig. 2B). Across the dataset, different subsets of precursor isotopes were isolated depending on their mass and charge (Extended Data Fig. 2A,B). We addressed these inconsistencies by modeling the quadrupole transmission function and using it to compute theoretical fragment isotope distributions. Doing so enabled deisotoping and recovery of total fragment intensities (Fig. 2C; Extended Data Fig. 2C). At the same time, some fragment annotations were unreliable, including cases with poorly fit isotope distributions or those outside the scan range, and were excluded from the loss function via masking (Fig. 2D; Extended Data Fig. 2D).

While harmonizing normalized collision energy (NCE) across runs, we observed that fragment intensities varied smoothly with NCE and that low-order splines accurately modeled this relationship (Extended Data Fig. 2E–H). This led us to design a different model architecture: rather than using NCE as an input to the neural network to predict intensities at a single energy, Altimeter predicts spline coefficients for each fragment. NCE is then supplied to the spline to evaluate intensity at arbitrary energies without additional inference (Fig. 2E; Extended Data Fig. 2I,3A). Because these predictions represent total fragment intensities, Pioneer can re-isotope library spectra to match scan-specific precursor isolation efficiencies (Extended Data Fig. 3B). This design improves speed and scalability. A spectral library needs to be predicted only once for a given search space—defined by the sequence database and digestion and modification settings—after which it can be reused across datasets.

**Fig 3:**
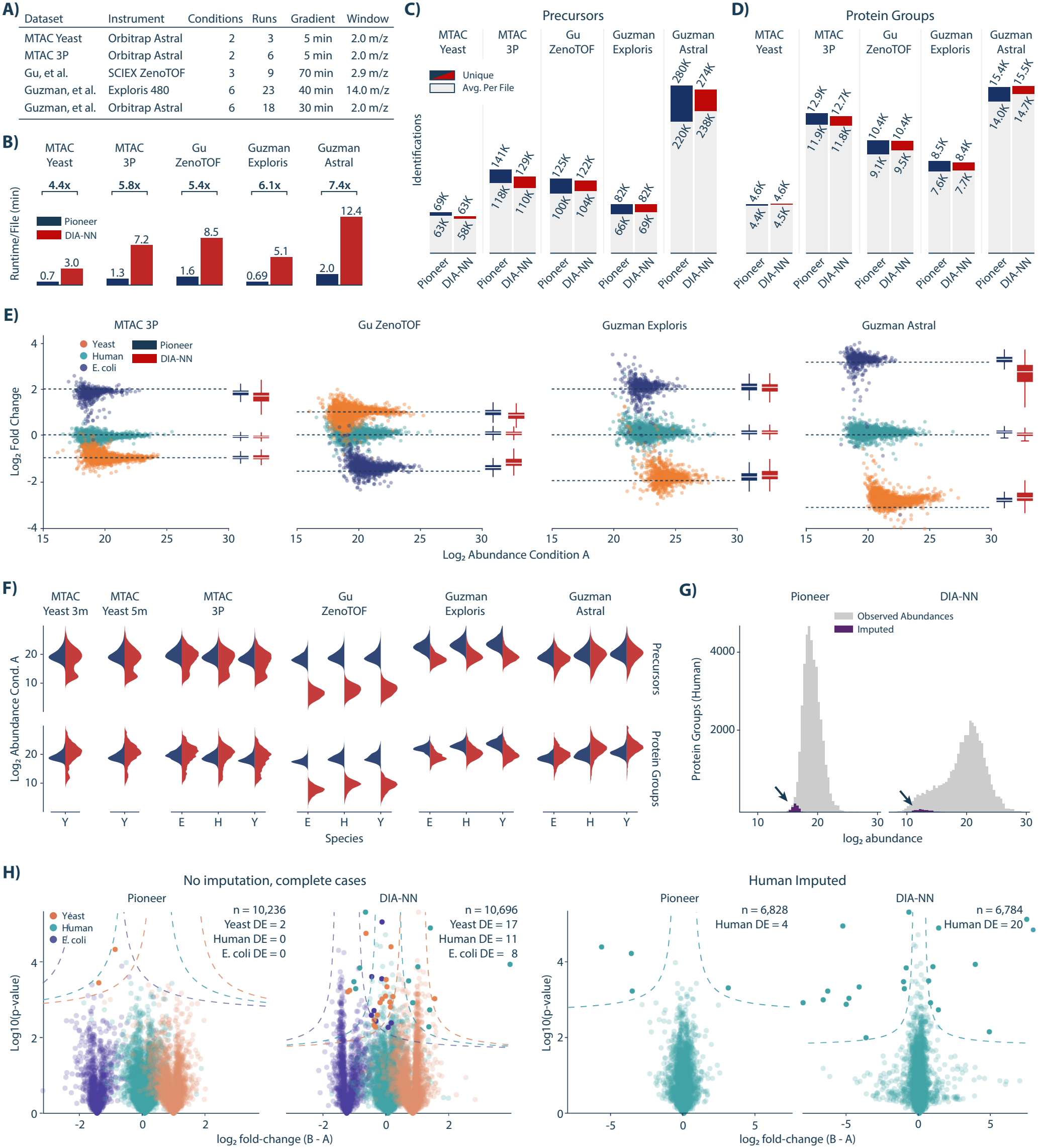
Pioneer performance on quantitative three-proteome benchmarks. **A)** Overview of the benchmark experiments. **B)** Analysis runtimes for Pioneer and DIA-NN v2.6.0. Times exclude library generation and file conversion for both tools. **C)** Unique and average precursor identifications. **D)** Unique and average protein group identifications. **E)** Log_2_ -fold change (conditions B vs A) versus log_2_ abundance for each protein group using Pioneer. The boxplots summarize Pioneer and DIA-NN results. **F)** Log_2_ abundance distributions. **G)** MTAC 3P example run showing human protein abundances and Perseus-imputed values with default parameters (width 0.3, downshift 1.8). Imputation was performed only when at least one condition had no missing values. **H)** Differential expression analysis (Perseus) for the MTAC 3P benchmark. For each species, differential expression was tested relative to its median log_2_ fold change using a two-sample t-test with a 5% FDR and s_0_ = 0.1. Left: no imputation, including all species; right: analysis restricted to human protein groups after imputing missing values.

To assess prediction accuracy, we evaluated Altimeter on a held-out test set comprising 10% of the data and achieved a median spectral angle of 0.9375 on the deisotoped spectra (Fig. 2F). Predicted fragment splines closely matched smoothed experimental trends across collision energies, demonstrating that the model captures the continuous dependence of fragmentation on NCE (Fig. 2G). Figure 2H shows a representative example in which a precursor spans two adjacent scans. Despite substantial differences in the observed fragment isotope peaks, Altimeter accurately predicts intensities in both spectra, accounts for a larger fraction of the MS2 signal than Prosit, and achieves higher Scribe scores with comparable spectral angles despite additional matching peaks [27, 39].

### 2.3 Pioneer excels in benchmark experiments

We next compared Pioneer v0.9.0 and DIA-NN v2.6.0 on replicate injections of a yeast proteome and on three-proteome benchmarks (Fig. 3A) [8, 40]. Pioneer completed its analyses 4.4-7.4x faster than DIA-NN (Fig. 3B) and scaled well with increasing CPU threads (Extended Data Fig. 4). Pioneer identified as many or more unique precursors in every benchmark by up to 10% and more unique protein groups in every experiment except for the Orbitrap Astral benchmark from Guzman et al. (Fig. 3C-D)[8]. In terms of average protein-group identifications per file, Pioneer was comparable with 4% fewer to 1% more than DIA-NN (Fig. 3D). Pioneer also more accurately recovered the expected fold changes, with lower variance and a larger dynamic range in the three proteome experiments (Fig. 3E).

**Fig 4:**
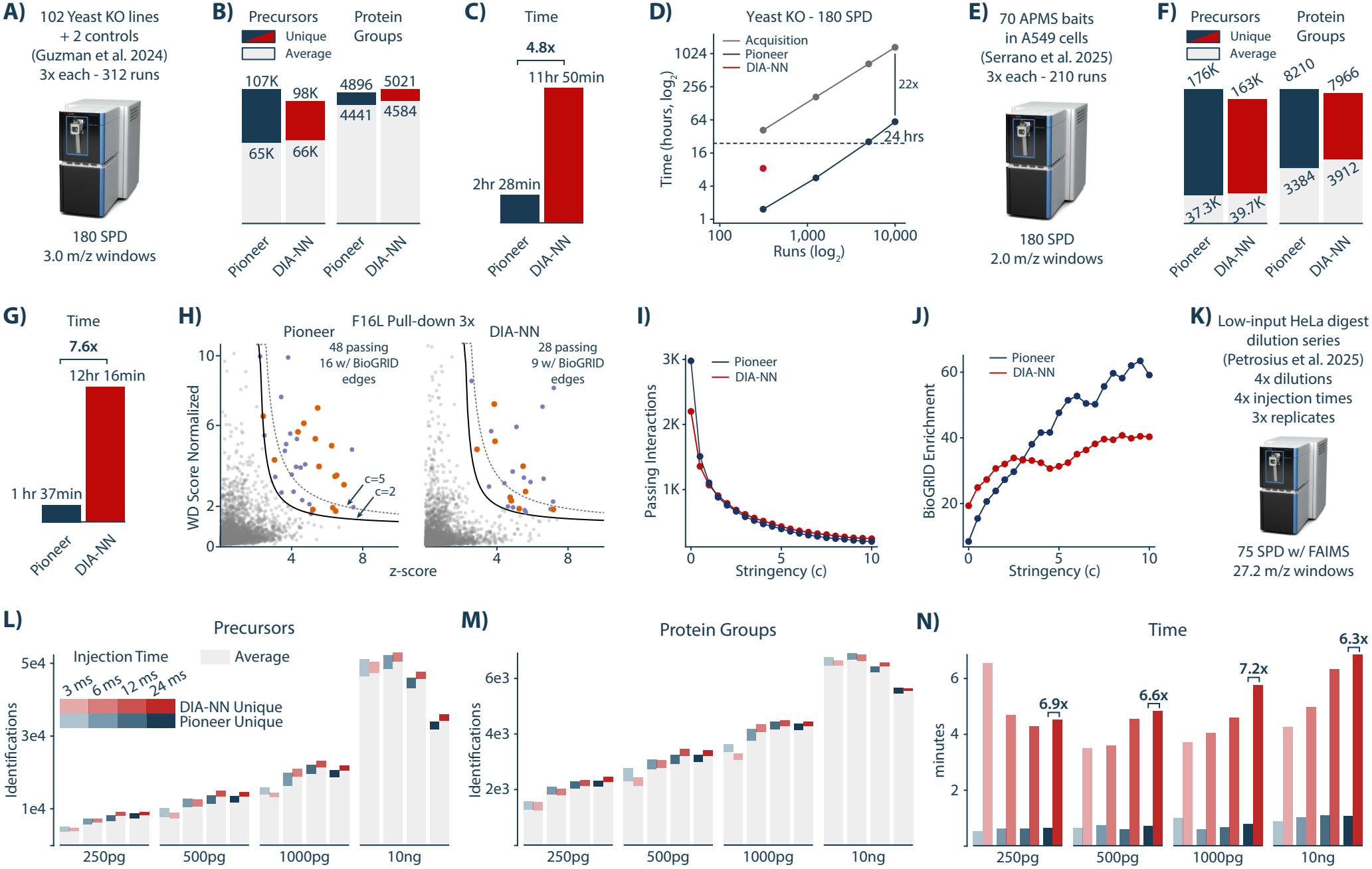
Pioneer performance across diverse experimental datasets. **A)** Schematic of the yeast knockout (KO) experiment from Guzman et al. [8]. **B)** Number of unique and average identifications for the yeast KO experiment. **C)** Runtime comparison for the yeast KO experiment. **D)** Runtime scaling with the number of runs. The number of runs was artificially increased using symbolic links to the original files, and the data were re-searched at 312, 1,248, 4,992 and 9,984 runs. The scaling analyses were limited to 16 threads and 64 GB of memory on a dedicated compute node with two Intel Xeon Gold 6548Y+ processors. Acquisition time is shown in grey, and DIA-NN and Pioneer runtimes in red and navy, respectively. DIA-NN ran out of memory before completing a 624-run analysis with the 64 GB memory limit. **E)** Schematic of the APMS experiment, in which vaccinia virus proteins were used as baits [13]. **F)** Number of unique and average identifications for the APMS experiment. **G)** Runtime comparison for the APMS experiment. **H)** WD-scoring analysis for an example bait, F16L. Because vaccinia bait proteins are absent from interaction databases, we assessed prey-prey interactions among co-purified human proteins. Y-axis: WD-scores normalized to 95th percentile. X-axis: prey abundance Z-score. Purple dots: proteins passing c ≥ 2 threshold. Orange dots: passing proteins with a prey partner also passing and present in the BioGRID physical multi-validated network [44]. **I)** Bait-prey pairs passing each stringency threshold. **J)** Enrichment of BioGRID-validated prey-prey interactions over a null model (100 permuted networks) at each stringency threshold. **K)** Schematic of the single-cell-equivalent experiment from Petrosius et al. [11]. **L)** Average and unique precursor identifications for the single-cell-equivalent experiment. **M)** Average and unique protein group identifications for the single-cell-equivalent experiment. **N)** Runtimes for the single-cell-equivalent experiment.

We used the MTAC yeast experiment to showcase the effects of including multiple fragment isotopes in the analysis. We re-searched the data but incorporated different numbers of fragment isotopes (Extended Data Fig. 5A). Addition of the M+1 isotopes (the default setting in Pioneer) improved precursor identifications by 5% but only marginally improved protein group identifications (Extended Data Fig. 5A). Further addition of the M+2 isotope improved neither precursor nor protein group identifications (Extended Data Fig. 5A). On the same experiment, Pioneer identified 19% more protein groups than AlphaDIA while completing the analysis 8.7 times faster [41] (Extended Data Fig. 5A,B).

**Fig 5:**
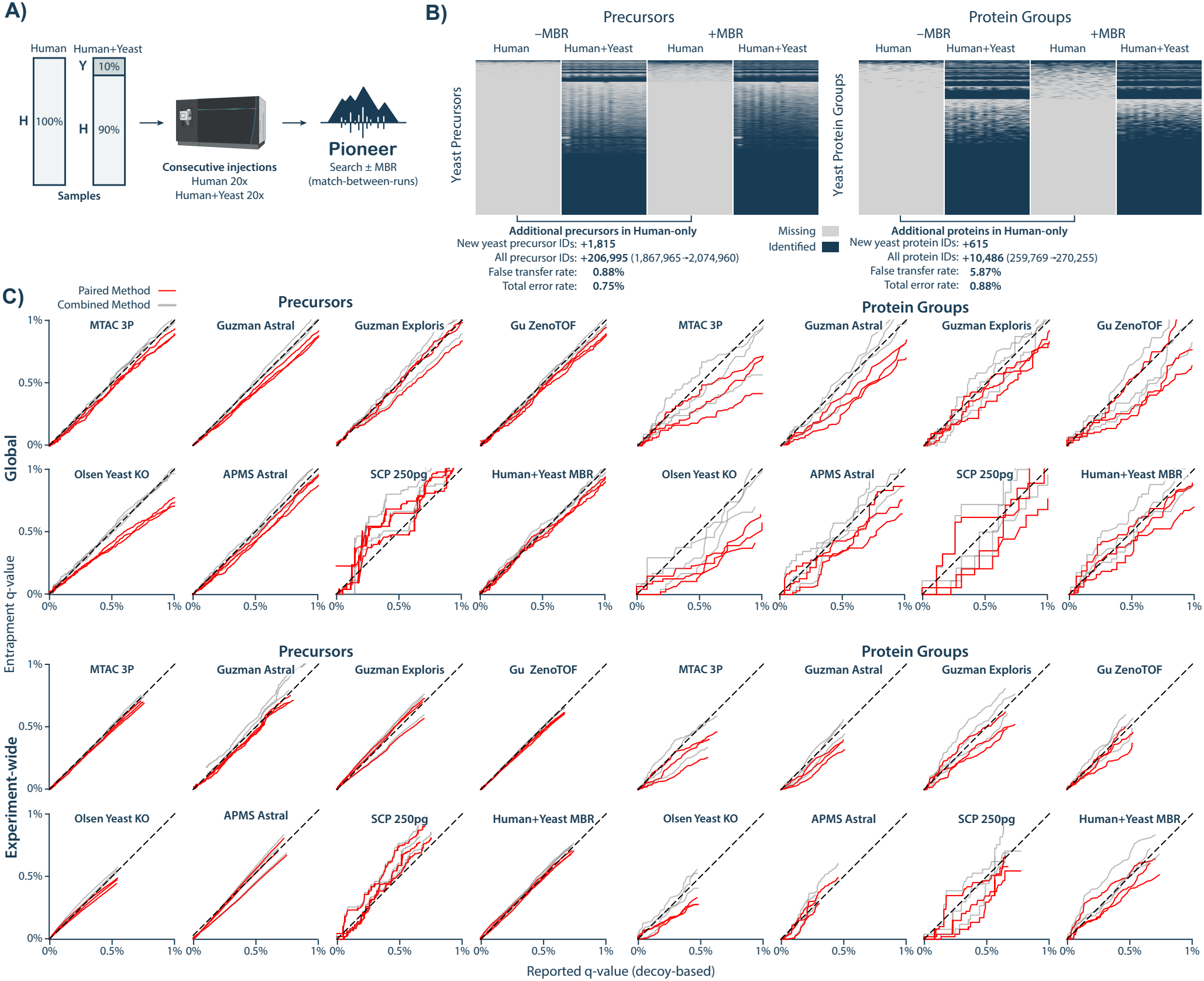
False-transfer rate evaluation and entrapment analyses. **A)** Experimental design for false-transfer rate (FTR) estimation: twenty injections of a tryptic human digest followed by twenty injections of a human-yeast mixture. Data were searched using Pioneer with and without match-between-runs (MBR). **B)** Yeast identifications in the human-only and the two-proteome runs with MBR on or off. Rows represent yeast precursors or protein groups and columns represent runs. Missing values are in grey and non-missing in navy. The false transfer rate and total error rate for precursors and protein groups are shown below the tables. **C)** Entrapment analyses for the three-proteome benchmarks (Fig. 3A), the yeast knockout, the APMS, the single-cell-equivalent experiments (Fig. 4A,E,K), and the two-proteome MBR experiment in (A). Identifications were required to pass 1% thresholds simultaneously on all four q-values (precursor and protein group, each global and experiment-wide). Entrapment analyses for the MTAC yeast benchmark and a further entrapment analysis of the single-cell-equivalent benchmark are shown in Extended Data Fig. 8B,C.

To evaluate the benefit of combining signal across isolation windows, we compared single-window and dual-window quantification in the MTAC yeast experiment. Dual-window quantification (default) combines precursor chromatograms from multiple isolation windows when a precursor’s isotopic envelope straddles each of these adjacent windows. We also tested whether increasing the time separation between adjacent windows improved quantification by re-acquiring the data with an alternating window method and analyzing both standard and alternating acquisitions with both quantification schemes (Extended Data Fig. 5C-D). The alternating window method was designed to maximize the time separation between quadrupole isolation windows within each cycle that are adjacent in their targeted m/z. The dual-window analysis method improved the average number of points measured across each chromatographic peak by nearly 1/3 (Extended Data Fig. 5F), while the alternating windows configuration did not improve quantification (Extended Data Fig. 5E). Owing to software constraints on the Orbitrap Astral, both the standard and alternating methods were approximate implementations (Extended Data Fig. 5G-K).

Protein abundance distributions are expected to be approximately log-normal [42]. In contrast, DIA-NN v2.6.0 produced heavy-tailed or bimodal distributions in some benchmarks, most prominently in the MTAC yeast and MTAC three-proteome datasets (Fig. 3F). To assess how these distributions affected imputation, we applied Perseus’s standard imputation method and found the imputed DIA-NN abundances frequently exceeded observed values (Fig. 3G) [43]. We therefore tested for false-positive differentially expressed proteins before and after imputation using a standard analysis pipeline in Perseus (Fig. 3H). In the MTAC three-proteome experiment, Pioneer reported no human false positives without imputation compared to 11 for DIA-NN, and likewise reported only 4 false positives with imputation compared to 20 for DIA-NN.

### 2.4 Pioneer supports large-scale and low-input datasets

Having established performance on controlled benchmarks of standard acquisition methods, we next evaluated Pioneer on three diverse experiments from the literature: a panel of 102 yeast knockouts in technical triplicate [8], a set of 70 affinity-purification mass spectrometry (APMS) experiments using vaccinia virus proteins as baits [13], likewise in triplicate, and lastly a series of single-cell-equivalent benchmarks acquired across an array of instrument parameters [11].

In the yeast knockout dataset (Fig. 4A), Pioneer identified nearly 10% more unique precursors than DIA-NN and identified within 2-3% as many unique and average protein group identifications, while completing the analysis 5-times faster (Fig. 4B,C). To assess consistency, we checked whether replicates from the same condition grouped together. We first imputed missing values to obtain a complete matrix for principal component analysis (PCA). Replicates from the same condition were indeed similar in PCA space (Extended Data Fig. 6A). The effects of imputation in our earlier benchmarks prompted us to test whether it also altered clustering. We therefore performed unsupervised agglomerative clustering before and after imputation, using 104 clusters to match the number of known conditions. Without imputation, all replicates of the same condition clustered together for 92% and 94% of conditions for Pioneer and DIA-NN, respectively (Extended Data Fig. 6B). Although imputation reduced these percentages, Pioneer better preserved replicate grouping (82% vs. 62% for DIA-NN) (Extended Data Fig. 6B).

We next examined how Pioneer’s runtime scales with the number of runs. We increased the yeast knockout experiment from 312 runs to 1,248, 4,992 and 9,984 runs using symbolic links to the original files. We searched these expanded datasets on a single compute node limited to 16 threads and 64 GB of memory (Fig. 4D). Pioneer analyzed all 9,984 runs in approximately 60 hours, 22 times faster than the projected acquisition time. DIA-NN ran out of memory before completing a 624-run analysis with the 64 GB memory limit. We performed a similar analysis on a 500 SPD HEK293T dataset acquired on an Orbitrap Astral Zoom [45], and likewise expanded the dataset from 550 to 2,200 runs (Extended Data Fig. 7). Pioneer analyzed all 2,200 runs in just under 24 hours, 4.6 times faster than acquisition. DIA-NN completed the original 550-run analysis only after its memory allocation was increased from 64 to 128 GB.

In the affinity-purification dataset (Fig. 4E), Pioneer identified 8% and 3% more unique precursors and protein groups respectively than DIA-NN, and completed its analysis 7.6-times faster (Fig. 4F,G). Pioneer identified 6% fewer average precursors and 15% fewer average protein groups per run (Fig. 4F). However, the APMS data enabled a biological validation of the identified protein groups, since we could look for physically validated prey-prey interactions among human proteins that were statistically associated with each given vaccinia virus bait protein (Fig. 4H). We found that from moderate to high stringency, Pioneer exhibited greater enrichment than DIA-NN of BioGRID physical, multi-validated, prey-prey interactions relative to permuted control networks (Fig. 4I,J).

Finally, Pioneer was evaluated on a single-cell-equivalent dilution series from 250 pg to 10 ng. Each dilution was analyzed with MS2 injection times of 3, 6, 12, and 24 milliseconds and ran in triplicate with a method length of approximately 16 minutes (Fig. 4K). Across these conditions Pioneer identified from 11% fewer to 15% more unique precursors than DIA-NN, and from 6% fewer to 14% more unique protein groups. Likewise, Pioneer identified from 16% fewer to 5% more average precursors than DIA-NN and from 9% fewer to 9% more average protein groups (Fig. 4L,M). Pioneer therefore had comparable identification depth with a 3.8 to 12 times faster analysis than DIA-NN. Moreover, Pioneer performed its analyses a median of 23 times faster than data acquisition (Fig. 4N).

### 2.5 Pioneer controls false-discovery and false-transfer rates

Lastly, we validated Pioneer’s false-transfer rate (FTR) and FDR control. Correct identifications can be transferred to runs in which they are absent. These errors can occur without introducing decoy or entrapment hits and may therefore escape FDR estimates. To estimate the FTR, we generated a two-proteome MBR dataset following a previously established strategy [46] (Fig. 5A). Yeast identifications were counted as false transfers if they appeared in the human-only runs only after MBR was enabled and were absent from those runs entirely with MBR off. The false-transfer count was divided by the gain in total identifications between the MBR-off and MBR-on analyses to obtain the FTR. The estimated FTR was 0.88% at the precursor level and 5.87% at the protein group level (Fig. 5B). Although the protein group FTR exceeded 1%, it only reflects errors in additional identifications. To assess the overall impact, we combined the decoy identification rate and false transfers relative to all identifications. The resulting total error rate was 0.75% at the precursor level and 0.88% at the protein group level.

Entrapment analyses across all datasets confirmed appropriate control of FDRs. The analyses were performed in triplicate, that is, with three separate entrapment libraries, where the shuffled entrapment sequences were randomly generated each time. Entrapment q-values closely tracked decoy-based q-values at the global and experiment-wide levels (Fig. 5C, Extended Data Fig. 8A-C). The paired entrapment method from Wen et al. showed no evidence of inflated false discovery rates [47]. The SCP analysis at 250 pg and 24 ms injection time was the only benchmark for which the average combined entrapment q-value was greater than the reported q-value. Despite the modest miscalibration, the estimate remained below 1%, supporting robust FDR control across all datasets.

## 3 Discussion

The Pioneer and Altimeter suite meets or exceeds state-of-the-art standards for peptide and protein group identification depth, quantitative accuracy, and FDR control for DIA proteomics. Pioneer furthermore completes a typical analysis four to eight times faster than previously possible and scales to 10,000 file searches on standard hardware, with no distributed computing required. Several principal innovations enable these capabilities. First, Pioneer uses a deep and curated list of statistics to discriminate target from decoy peptides in a multi-stage search. Second, to rapidly reduce the list of candidate precursors, Pioneer implements a fragment index that accounts for relative fragment abundances. Third, Pioneer uses a fast and robust linear model to regress empirical spectra onto library spectra. The model coefficients feed into both quantification and calculation of spectral similarity features. Fourth, Altimeter spectral libraries model relative fragment abundance as a continuous function of a collision energy parameter. Lastly, Pioneer estimates run-specific quadrupole transmission functions from the data and then uses these models to re-isotope Altimeter library spectra for each individual MS2 scan.

Several studies have noted that DIA MS2 spectra differ from DDA MS2 spectra due to increased co-fragmentation, and the absence of charge-dependent collision-energy optimization [48–50]. Our results further show that differences in precursor isotope isolation alter each fragment’s isotopic envelope and contribute substantially to this divergence. This has consequences for both library construction and quantification. Empirical libraries are implicitly tied to the isolation windows under which they were generated. For a given precursor, a library may reflect only a single window, leading to preferential identification in that window and misquantification in others. Changes to isolation windows within a dataset would further exacerbate these effects and introduce systematic quantitative biases. Pioneer corrects for these effects, making quantification robust to differences in isolation windows.

In practice, effective isolation can change even when the acquisition method is held constant. Drift in quadrupole transmission over time, instrument recalibration or maintenance, and acquisition across multiple instruments all alter precursor isolation. The same factors also introduce differences in collision energy. Transfer learning and fine-tuning can partially absorb such shifts, but they are computationally expensive and typically applied globally rather than separately for each file. In contrast, Altimeter decouples prediction from acquisition by using peptide sequence and charge as its core inputs. Collision energy and isolation effects are applied during analysis to enable per-run adaptation without retraining. This architectural separation positions Altimeter as a potential foundation model, where a shared sequence- and charge-based representation could support distinct downstream decoders for different fragmentation methods or collisional cross-section prediction.

In conclusion the analyses presented here establish Pioneer and Altimeter as a tested, open-source and cross-platform solution for rapid analysis of DIA proteomics data that combines deep proteome coverage with quantitative accuracy and strict control of false discovery rates. Pioneer is designed to scale to large datasets with only modest memory and CPU requirements found on standard, personal computers. This software suite provides a practical foundation for modern large-scale DIA studies, and includes a graphical user interface (GUI) that facilitates practical analysis end-to-end.

## 4 Methods

### 4.1 MTAC Benchmark Experiments

Three-proteome and yeast benchmarks (Fig. 3) were performed at the Mass Spectrometry Technology Access Center at the McDonnell Genome Institute (MTAC@MGI) at Washington University School of Medicine. Commercial protein digest standards were obtained for *Homo sapiens* (Pierce HeLa Protein Digest Standard, Thermo Scientific, Cat. No. 88329, Lot UF286759), *Saccharomyces cerevisiae* strain BY4741 (Pierce Yeast Digest Standard, Thermo Scientific, Cat. No. A47951, Lot AB407858), and *Escherichia coli* (MassPREP E. coli Digest Standard, Waters Corporation, Part No. 186003196, Lot W07052404). Two three-proteome mixtures were prepared by combining the digests at the following mass ratios: Condition A contained 50% human, 20% yeast, and 30% *E. coli*, while Condition B contained 50% human, 40% yeast, and 10% *E. coli*. Stock solutions were diluted to 80 ng/µL in 98% water, 2% acetonitrile, and 0.1% formic acid. A 5 µL injection (400 ng on column) was used for all samples.

Samples were analyzed on an Orbitrap Astral mass spectrometer (Thermo Scientific) coupled to a Vanquish Neo LC system with a µPAC Neo 5.5 cm column (75 µm ID) maintained at 50°C. Mobile phase A was water with 0.1% formic acid; mobile phase B was 80% acetonitrile with 0.1% formic acid. Peptides were separated at 1.5 µL/min using a 5-minute active gradient from 12.6% to 45% B. MS1 scans were acquired in the Orbitrap at 240,000 resolution (380–980 m/z, 5 ms max IT, 500% AGC). DIA scans were acquired in the Astral analyzer using 280 windows of 2 Th isolation width spanning 392–948 m/z (150–2000 m/z scan range, 25% NCE, 5 ms max IT, 0.6 s cycle time). Three technical replicates were acquired for each condition. A yeast-only digest was analyzed in triplicate using both 3-minute and 5-minute active gradients.

### 4.2 Alternating DIA scheme

To design an alternating isolation window method compatible with the Orbitrap Astral, we first acquired data using sequential m/z windows (standard method) and extracted the realized MS2 scan order from the raw file. Two sub-cycles were detected, corresponding to quadrupole rod polarity groups. We then constructed a revised method in which isolation windows were reordered to maximize temporal separation between m/z-adjacent windows while preserving consecutive acquisition within each polarity block. This produced the most alternating schedule achievable without violating polarity grouping or triggering scan reordering.

In the standard method, approximately half of m/z-adjacent isolation windows (2 m/z apart in center mass) were acquired consecutively, and the remainder were separated by approximately one-half of a scan cycle (*∼*140 scans). In the alternating method, no m/z-adjacent windows were acquired consecutively; separations were distributed across the cycle, with approximately one quarter separated by one quarter of a cycle (*∼*70 scans), one half by one half of a cycle (*∼*140 scans), and one quarter by three quarters of a cycle (*∼*210 scans).

### 4.3 Spectral Library Generation

Spectral libraries were generated by tryptic *in silico* digestion of the UniProt reference proteomes for *Homo sapiens* (UP000005640), *Saccharomyces cerevisiae* (UP000002311), and *Escherichia coli* strain K12 (UP000000625), using canonical sequences without isoforms. Parameters common to all tools included cleavage before proline, 1 missed cleavage, carbamidomethylation of cysteine as a fixed modification, 1 variably oxidized methionine, 7-30 AA peptide length, and precursor charges 1–4. For all datasets, the instrument scan m/z ranges were fully covered by their corresponding libraries. Altimeter library precursor and fragment m/z ranges were automatically detected and trimmed to match the experiment files. DIA-NN libraries were generated using its built-in model with the default precursor m/z range, 300–1800, and fragment m/z range, 200–1800. AlphaDIA libraries were generated using AlphaPeptDeep [25] with a precursor charge range of 2-4, a precursor m/z range from 359–1034, a fragment m/z range of 150–2000, b/y ions with maximum charge 2, and NCE 26 with the Astral instrument model.

### 4.4 DIA Software Analyses

Pioneer analyses were performed using version 0.9.0 with default parameters unless otherwise indicated. Pioneer version 0.12.0 was used for the analyses in Fig. 4D and Extended Data Fig. 7. DIA-NN analyses were performed using version 2.6.0 with default parameters, except protein inference was set to “Protein Names”. AlphaDIA analyses were performed using version 2.1.3 with default parameters; MBR was enabled, FDR was controlled at the protein level using heuristic inference, and quantification used directLFQ normalization. For all three tools, results were filtered to a 1% FDR threshold. Run-to-run normalization was disabled for Pioneer and DIA-NN because several datasets (e.g., three-proteome mixtures and APMS) violate the assumption of global proteome stability across runs, and identification benchmarks are unaffected by normalization. Unless otherwise stated analyses were performed on a computer running Windows 10 with a 3.0 GHz 18-core Intel i9-10980XE CPU, 64 GB 2666 MHz DDR4 RAM, using 24 threads.

### 4.5 Statistics and Imputation

For the differential expression analysis of the MTAC three-proteome experiment (Fig. 3G-H), imputation and statistical testing were carried out in Perseus [43]. Missing values were imputed by drawing from a normal distribution with a width of 0.3 standard deviations and a downshift of 1.8 standard deviations from the mean of the observed abundance distribution [51]. Imputation was restricted to protein groups for which at least one condition had no missing values. Differential expression was assessed in Perseus using a two-sample *t*-test (*s*_0_ = 0.1, 5% FDR). For the human-only differential expression analysis (Fig. 3H), we filtered out non-human protein-groups before imputation.

### 4.6 Protein-Protein Interaction Scoring and Evaluation

To validate protein-protein interactions from APMS data [13] (Fig. 4H-J), we applied WD-scoring as described by Sowa et al. and Huttlin et al. [52, 53]. WD-scores were normalized to the 95th percentile across all bait-prey pairs. We applied a combined WD-score and Z-score threshold to identify high-confidence interactions [13]. For each prey protein, we calculated the Z-score as 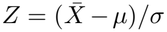 , where 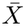 is the mean prey abundance for the given bait, and *µ* and *σ* are the global mean and standard deviation of the prey’s abundance across all samples. Prey proteins were considered interactors if they satisfied:

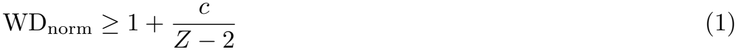

where *c* is a stringency parameter. Prey proteins with *Z≤* 2 were excluded. We evaluated performance across stringency thresholds from *c* = 0 to *c* = 10. Missing values were not imputed; mean abundances were calculated from detected replicates only.

To assess biological validity, we identified putative prey-prey pairs found in the BioGRID physical multi-validated interaction network (version 5.0.252) [44]. Because vaccinia virus proteins served as baits and were absent from BioGRID, we restricted our analysis to prey-prey interactions among human proteins. We constructed 100 permuted networks by filtering BioGRID to human-to-human interactions and randomly shuffling the pairings. At a given stringency parameter, c, we counted prey-prey pairs where both members passed the scoring threshold. Enrichment was calculated as the ratio of BioGRID-validated pairs observed in the true network to the mean number observed across the permuted networks.

### 4.7 Entrapment Analysis

To evaluate false discovery rate control, we performed entrapment analysis following Wen et al. [47]. We generated target peptide sequences by *in silico* tryptic digestion of the protein FASTA database and created decoy sequences by shuffling target peptides while keeping the C-terminal amino acid fixed, re-shuffling up to 20 times if needed to avoid duplicates. We then applied the same shuffling procedure to these original target and decoy peptide sequences to generate paired entrapment peptides with an entrapment ratio of *r* = 1. During library construction, each target precursor received an entrapment pair ID linking it to its corresponding entrapment precursor.

We independently generated three replicate spectral libraries using this procedure to assess variability in the entrapment estimates, then searched DIA data against each library at both 1% and 5% q-value thresholds. We estimated the false discovery proportion using both the combined and paired methods [47].

For protein group-level analysis, the paired method required a matching strategy to account for differences in protein group composition between target/decoy and entrapment identifications. Because protein inference can assign different sets of protein accessions to target versus entrapment peptides, direct one-to-one pairing is not always possible. For each entrapment protein group, we therefore matched it to the target protein group sharing the maximum number of protein accessions.

### 4.8 False Transfer Rate Estimation

To estimate the false transfer rate (FTR) of match-between-runs, we prepared two sample types: HeLa digest alone and HeLa digest spiked with yeast digest at a 9:1 mass ratio. Commercial protein digest standards were obtained from Thermo Scientific Pierce: *Homo sapiens* (Cat. No. 88329) and *Saccharomyces cerevisiae* (Cat. No. A47951). Stock solutions were prepared at 200 ng/µL and a 1 µL injection (200 ng on column) was used for all samples.

Samples were analyzed on an Orbitrap Eclipse mass spectrometer (Thermo Scientific) coupled to an UltiMate 3000 RSLCnano system in trap-elute configuration. Peptides were trapped on a Pharmafluidics µPAC trap column and separated on a µPAC Neo 50 cm column at 300 nL/min using a 58-minute active gradient from 12% to 40% B (12% to 20% B over 42 min, then 20% to 40% B over 16 min). Mobile phase A was water with 0.1% formic acid; mobile phase B was 100% acetonitrile with 0.1% formic acid. MS1 scans were acquired in the Orbitrap at 30,000 resolution (390–1010 m/z, 54 ms max IT, 100% AGC, 30% RF lens). DIA scans were acquired in the Orbitrap at 30,000 resolution using 59 windows of 10 m/z isolation width with 1 m/z overlap, spanning 400–1000 m/z (145–1450 m/z scan range, 30% NCE, 54 ms max IT, 1000% AGC, 3 s cycle time).

We acquired 20 human-only replicates followed by 20 HeLa+yeast replicates and searched together with MBR enabled and disabled. Yeast peptides are absent from human-only samples. Therefore, we considered a false transfer any yeast precursor or protein group identification in any of the human only runs with MBR turned on that was not identified in at least one file in the human-only runs with MBR turned off. We divided the number of false transfers by the total gain in identifications between the MBR on and MBR off analyses. We calculated the total error rate by first taking the sum of false transfers and total decoy identifications and then dividing that sum by the total number of identifications.

## Data Availability

All mass spectrometry datasets used or generated in this study are publicly available via one of the PRIDE [54], MassIVE, or jPOST repositories [55].

Previously published datasets reanalyzed in this work include the ProteomeTools synthetic peptide dataset (PXD021013, PXD010595, PXD004732, PXD006832); three-proteome benchmark datasets acquired on Orbitrap Astral and Orbitrap Exploris instruments from Guzman et al. (PXD046444); the Guzman et al. yeast knock-out benchmark dataset (PXD046386); the Guzman et al. Orbitrap Astral Zoom 500 SPD HEK293T dataset (PXD071972); the affinity-purification mass spectrometry (APMS) dataset (MSV000096304); the SCIEX three-proteome benchmark dataset (JPST002949); and the single-cell-equivalent proteomics dataset (MSV000095333).

Datasets generated in this study and used in the main figures have also been deposited in PRIDE, including the two-proteome match-between-runs benchmark dataset (PXD074320), MTAC yeast benchmark datasets acquired with standard and alternating DIA acquisition schemes (PXD074310), and the MTAC three-proteome benchmark dataset (PXD074310).

## Code Availability

All software developed in this study is open source. Pioneer is available at https://github.com/nwamsley1/Pioneer.jl and is released under the GNU Affero General Public License v3 (AGPL-3.0). The Altimeter code is available at https://github.com/GoldfarbLab/Altimeter and is released under the MIT License. Altimeter is also deployed as a hosted inference service via Koina at https://koina.wilhelmlab.org/. Processed training data used for Altimeter model development are publicly available via Zenodo (DOI: 10.5281/zenodo.15875053).

## Acknowledgments

We thank the Mass Spectrometry Technology Access Center (MTAC) at Washington University in St. Louis for access to instrumentation and support for the Orbitrap Astral experiments. We also thank Ludwig Lautenbacher for his assistance in deploying Altimeter on the Koina inference server. M.B.M. was supported by NIH grant R01CA290809.

## Author contributions

N.T.W. developed Pioneer. E.M.W. performed mass spectrometry experiments and software testing. M.B.M. contributed to project oversight and mentorship. D.G. developed Altimeter and supervised the overall project.

## Notes

### Competing Interest Statement

The authors have declared no competing interest.

### Summary of Updates

Revisions to abstract and addition of Figure 4D) and Extended Data Figure 7. Results and Discussion were updated to discuss the new figures.

