## Supplementary Information for "Pioneer and Altimeter: Rapid and Scalable Analysis of DIA Proteomics Data Optimized for Narrow Isolation Windows"

### 1 Supplementary Methods

This section is organized to mirror the conceptual workflow of Pioneer. We begin with the construction and training of Altimeter, including data curation, fragment annotation, harmonization, model architecture, and deployment. We then describe Pioneer's raw file conversion, spectral library generation, and the intensity-aware fragment index used for rapid MS2 candidate precursor identification. The subsequent sections detail the Pioneer analysis pipeline itself, following the steps outlined in Extended Data Fig. 1: parameter tuning, the main search, spectral deconvolution, match-between-runs (MBR), semi-supervised model training, false-discovery and false-transfer rate control, chromatogram integration, and protein inference and quantification. Finally, we provide a mathematical treatment of fragment isotope correction and estimation of the quadrupole transmission function. Together, these sections describe the complete statistical and computational framework that underlies Pioneer and Altimeter.

#### 1.1 Altimeter Training Data

The training dataset for Altimeter was derived from the ProteomeTools project and was re-processed in-house. Raw files were obtained from PRIDE: PXD021013 (HLA and non-tryptic peptides), PXD010595 and PXD004732 (tryptic peptides), and PXD006832 (PROCAL calibration).

##### 1.1.1 Database Searching

Raw files containing HCD data were converted to mzML using ThermoRawFileParser v1.4.4 and searched with sage v0.14.7 against the human UniProt reference proteome (release 2024-06-04; canonical isoforms only), the PROCAL calibration peptides (PXD006832), and pool-specific FASTA files generated for each raw file. These pool-specific FASTAs contained both the intended synthetic peptide sequences and potential synthesis error variants, specifically double C-terminal residues, single amino acid deletions, and early termination products. Searches were performed with default parameters, with the following constraints: precursor and fragment tolerance $\pm10$ ppm, fully enzymatic specificity with up to three missed cleavages, peptide lengths 5–60 amino acids, peptide masses 500–8,000 Da, and precursor charge states +1 to +8. Carbamidomethylation of cysteine was set as a static modification, and oxidation of methionine as a variable modification. Spectrum-level FDR was controlled at 1% using a target–decoy approach.

##### 1.1.2 Spectrum Filtering

After database searching, only spectra acquired using higher-energy collisional dissociation (HCD) in the Orbitrap analyzer were retained. Additional quality control steps were applied to ensure that only high-confidence spectra were used for training. Precursor isolation purity was calculated from MS1 signals. It was required that the isolation window be centered on one of the isotopes of the identified precursor and also that the set of precursor isotopes together contribute at least 90% of the total signal within the isolation window. The MS1 isotope distribution also had to agree with the theoretical precursor distribution, with a cosine similarity $\geq$0.95. Peptide-spectrum matches were further filtered to those with a sage hyperscore $\geq$30. Finally, retention times were aligned to Chronologer predictions using spline fitting, and spectra were excluded if their observed retention times deviated by more than three standard deviations from the predicted values.

##### 1.1.3 Fragment Annotation

All spectra were initially annotated for b- and y-type ions, precursor ions, and immonium ions, including their isotopes. Annotation was performed with a 20 ppm tolerance. After annotation, ions were retained only if they were observed in at least 2% of the spectra in which they were theoretically detectable given the sequence, charge, and m/z scan range. This filtered set defined the ion dictionary used for model training.

Fragment m/z values were corrected on a per-spectrum basis by fitting the ppm error as a function of fragment m/z using the RANSAC algorithm, a robust regression method. Following this correction, annotation was repeated to recover ions that may have been missed during the first pass.

##### 1.1.4 Quadrupole Transmission Fit and Deisotoping

To identify fragment isotope clusters that were inconsistent with theoretical expectations, possibly due to interference, we fit the quadrupole transmission function. For each spectrum, precursor isotope contributions were estimated using the approach described in section 1.14.1, and ratios were computed relative to the theoretical precursor isotope distribution, assuming that the center isotope was isolated at 100%. Spectra from raw files that shared the same quadrupole calibration date were then jointly fit with an asymmetric generalized bell function, yielding a calibration-wide transmission profile (Extended Data Fig. 2C).

Fragment ions were then deisotoped by collapsing isotope clusters into single monoisotopic entries representing the total fragment intensity. When isotope clusters overlapped in m/z, their intensities were deconvolved according to the theoretical contributions of each overlapping isotope, and the resulting abundances were summed. The observed isotope distribution for each fragment was then compared to the theoretical distribution derived from the transmission profile, and any cluster with a cosine similarity below 0.9 was flagged as inconsistent.

##### 1.1.5 Fragment Masking

Each annotated fragment ion was assigned a mask indicating how it should be handled during training (Extended Data Fig. 2D). Five cases were defined:

1. Unmasked — the fragment passed all checks.
2. Outside m/z scan range — the fragment lay outside the MS2 m/z scan range.
3. High mass error — fragments with a mass error $\geq$ 15 ppm.
4. Inconsistent isotope distribution — fragments that failed the isotope consistency check.
5. Ambiguous annotation — fragments that could not be uniquely assigned because multiple ions shared the same m/z.

##### 1.1.6 NCE Alignment

Reported normalized collision energies (NCEs) were aligned to the PROCAL Lumos scale for training. The PROCAL data, part of the ProteomeTools project, had been collected on both the Fusion Lumos and QE instruments and were reprocessed in-house. For each spectrum, intensities were normalized by dividing by the total annotated intensity, and the median normalized intensity across replicates was taken for each fragment–NCE combination. On the QE, spectra were collected at 15 NCE settings, and fragment splines were fit to these data to provide a dense reference across the NCE range (Extended Data Fig. 2E).

To align Lumos PROCAL data to this reference, candidate NCE offsets were tested in 0.1-unit increments, and spectral angle was used to compare observed spectra with those predicted from the QE splines. For each precursor, the aligned NCE was defined as the median value that gave the best agreement. These aligned NCEs were then used to fit a mapping spline between QE and Lumos values (Extended Data Fig. 2F). For experimental runs, reported NCEs were first aligned to the QE fragment splines and then converted to the PROCAL Lumos scale using this mapping. The median result across precursors was taken as the aligned NCE for that file, and the difference between the aligned and reported values was defined as the NCE offset:

$$\mathrm{NCE}_{\mathrm{aligned}} = \mathrm{NCE}_{\mathrm{reported}} + \mathrm{NCE}_{\mathrm{offset}} \tag{1}$$

Offsets were then smoothed over time with a 12-hour rolling window, applied separately to each reported NCE across all raw files collected under the same collision-energy calibration date (Extended Data Fig. 2G,H).

##### 1.1.7 Altimeter Model Architecture

Altimeter is a transformer-based model built on the architecture used by UniSpec [1] (Extended Data Fig. 2I). The input consists of the peptide sequence and precursor charge. Sequences are one-hot encoded and projected into a 256-dimensional embedding space; charge is embedded separately and concatenated with the sequence representation before being passed through nine transformer blocks with a dropout rate of 0.1. The model contains 11.8 million trainable parameters and was implemented in PyTorch. The output is a set of cubic B-spline coefficients for each fragment ion in the ion dictionary. Four coefficients are predicted per fragment, along with eight knot positions that are learned for the entire model. Given an aligned NCE, splines are evaluated to produce a predicted monoisotopic, total-intensity spectrum.

##### 1.1.8 Training

Altimeter was trained with the Adam optimizer (learning rate $3 \times 10^{-4}$, exponential decay 0.85) using a batch size of 100. Data were split 70/20/10 into training, validation, and test sets, with uniform sampling.

The objective was a masked spectral-angle loss on the predicted total-intensity spectra. Fragments outside the MS2 $m/z$ range were excluded, while those with high mass error, inconsistent isotope distributions, or ambiguous annotations contributed only as upper bounds: predictions below the observed intensity were ignored, and over-predictions were penalized in proportion to the excess. Predictions below a spectrum's smallest detected signal were also excluded from the loss. Each spectrum was weighted by the square root of precursor isolation purity multiplied by rawOvFtT to prioritize high-quality, abundant spectra.

Spectral angle was computed from the cosine similarity ($cs$) between predicted and observed spectra. Cosine values were clamped to the range $[-(1-\epsilon), (1-\epsilon)]$ with $\epsilon = 1\times10^{-5}$ to ensure numerical stability, then converted as

$$SA = -\left(1 - 2 \cdot \frac{\arccos(cs)}{\pi}\right).$$

Training ran for up to 100 epochs with validation tested after each epoch. Early stopping with a patience of five epochs selected the model from epoch 30 as optimal (Extended Data Fig. 2J).

##### 1.1.9 Deployment

Four inference variants are deployed via the Koina framework:

1. Altimeter_2024_intensities: produces the predicted total-intensity spectrum (splines evaluated at requested NCE) and returns fragment intensities.
2. Altimeter_2024_isotopes: returns re-isotoped spectra so that fragment isotope distributions (given precursor isotope transmission) can be reconstructed.
3. Altimeter_2024_splines: returns the raw spline coefficients for fragment ions (without evaluating them at a specific NCE) so downstream users can evaluate them as needed.
4. Altimeter_2024_splines_index: similar to splines variant but uses integer fragment ion indices (not string annotation names) for faster performance and lighter I/O.

#### 1.2 Cross-Platform Mass Spectrometry File Conversion

Pioneer reads mass spectrometry data in the Apache Arrow IPC format rather than directly from the vendor formats. The PioneerConverter tool converts Thermo Scientific Raw (.raw) files to the IPC format.

PioneerConverter is distributed both as a standalone tool (https://github.com/nwamsley1/PioneerConverter) and packaged with the Pioneer installer. It uses the Thermo Scientific RawFileReader to read the .RAW files and is cross-platform. In addition, Pioneer supports the conversion of .mzML files to the same Arrow format.

#### 1.3 Spectral Library Sequence Generation

Pioneer constructs target-decoy sequence libraries as inputs to Altimeter and Chronologer [2] for *in silico* spectrum and retention time prediction. Given a list of FASTA-formatted protein sequences, Pioneer digests these according to a user-defined enzymatic cleavage rule. After the initial digestion, Pioneer generates decoy sequences by either reversing or shuffling the target peptide sequences according to a user-defined parameter. In either case the C-terminal amino acid remains fixed in each sequence, and the mass distribution of the decoy sequences matches that of the targets [3]. For internal testing, Pioneer optionally adds entrapment sequences to evaluate its false discovery rate estimation according to Wen et al. [4]. Pioneer generates the entrapment targets by randomly shuffling each original target sequence while keeping the C-terminal amino acid fixed. Because reversing a given target to form both a decoy and an entrapment sequence would yield identical sequences, Pioneer does not permit both the decoy and entrapment sets to be generated by reversal; if one set is generated by reversal, the other is generated by shuffling. Treating isoleucine and leucine as identical, Pioneer shuffles until the peptide is unique across all targets, decoys, and entrapment peptides, or until 20 consecutive attempts fail, in which case the decoy or entrapment is skipped.

#### 1.4 Intensity-Aware Fragment Index Search

Both MSFragger and Sage use a fragment-index search to efficiently look up precursors from a digested sequence database for which several theoretical fragments match the experimental MS2 spectra [5, 6]. Pioneer implements a modified version of this data structure that accounts for spectral library fragment intensities. For each peak in an MS2 spectrum, Pioneer performs an index query that records all library ions that satisfy the retention time, fragment m/z, and precursor m/z constraints. The fragment-index construction and query process are as follows.

##### 1.4.1 Index Construction

The fragment index is constructed as follows.

1. Low-specificity fragments ($y_1$, $y_2$, $y_3$, $b_1$, $b_2$), isotope peaks, and precursor ions are removed.
2. For each precursor, the top eight fragments are retained. Rankings are determined by area under the relative intensity versus NCE curve. Each retained fragment is assigned a one-hot bitmask. The $r$-th ranked fragment ($r = 1, \ldots, 8$) is given the value $2^{r-1}$.
3. Each retained fragment is stored as a tuple

$$I = (P_k, S_k)_{k=1}^{m},$$

where $P_k$ is a partition-local precursor identifier (see below) and $S_k$ is the fragment rank bitmask.

The fragment index is organized in a three-level hierarchy—precursor m/z partitions, which each contain retention time bins, which each contain fragment m/z bins.

1. **Precursor m/z partitions.** The library is divided into contiguous precursor m/z intervals (default width 5 Da). Precursor identifiers are remapped to 16-bit indices within each partition, so any partition that would contain more than 65,535 precursors is recursively split.
2. **Retention time bins.** Within each partition, fragments are partitioned into non-overlapping retention time intervals

$$[RT_{lo,k}, RT_{hi,k}].$$

Each retention time bin stores:

- Its lower and upper bounds,
- Pointers to the first and last fragment m/z bins contained within the interval.

3. **Fragment m/z bins.** Within each retention time bin, fragments are further partitioned into non-overlapping fragment m/z intervals

$$[M_{lo,j}, M_{hi,j}].$$

Each m/z bin stores:

- Its lower and upper bounds,
- Pointers to the first and last entries in $I$ corresponding to fragments in the bin.

The precursor m/z, retention time, and fragment m/z bins are sorted in ascending order respectively.

##### 1.4.2 Score Counter

During fragment-index search, Pioneer maintains a counter that accumulates evidence for candidate precursors within a single MS2 scan. The counter consists of:

- An array $V$ listing the precursor IDs matched in the current scan, in the order they are first encountered,
- An array $W$, indexed by precursor ID, holding each precursor's accumulated fragment bitmask,
- An integer $n$ tracking the number of unique precursors encountered.

When a library fragment matches a peak, its rank bitmask (Section 1.4.1) is combined into the matched precursor's entry of $W$ by bitwise OR; the first time a precursor is matched, its ID is written to the $n + 1$ position of $V$, and $n$ is incremented. After all peaks are processed, $W$ holds, for each matched precursor, the bitwise OR of the rank bitmasks of its observed fragments, encoding which of its top-ranked fragments were detected.

##### 1.4.3 MS2 Scan Query

For each MS2 scan, Pioneer performs a fragment-index query constrained by the scan precursor m/z window $[mz_{min}, mz_{max}]$ and retention time window $[rt_{min}, rt_{max}]$ (library retention time units), producing candidate precursors for downstream spectral deconvolution. The query proceeds as follows.

1. Select the index partitions whose precursor m/z ranges overlap with $[mz_{min}, mz_{max}]$, and initialize the counter by setting $n = 0$.
2. Within each partition, Pioneer uses a binary search to find the first matching retention time sub-partition, and then sequentially processes retention time bins so long as they are within the retention time tolerance of the given MS2 scan.
3. For each retention time bin and each peak in the spectrum:
   - Determine the peak's fragment m/z tolerance interval $[x_{lo}, x_{hi}]$ from the mass-error model.
   - Locate the fragment m/z bins overlapping $[x_{lo}, x_{hi}]$ using a hybrid binary-then-linear search, seeded from the previous peak's bin and finished with a vectorized (SIMD) scan.
   - For each fragment $(P_k, S_k)$ in the matching bins, combine its rank bitmask $S_k$ into precursor $P_k$'s counter entry by bitwise OR.
4. After all peaks are processed, look up each encountered precursor's accumulated bitmask in the pass/fail table learned during bitmask calibration (Section 1.4.4); precursors with a passing bitmask are emitted as candidate matches for the scan. Precise precursor m/z and retention-time filtering are deferred to the spectral deconvolution step.
5. Reset the counter by clearing the accumulated bitmasks of the first $n$ precursors and setting $n = 0$.

##### 1.4.4 Bitmask Calibration

A precursor's accumulated bitmask (Section 1.4.2) is one of $2^8 = 256$ patterns that records which of its top eight library fragments were observed in a scan. Before the main search, Pioneer learns for each file a 256-entry pass/fail lookup table that decides, for each pattern, whether precursors matching it should be retained as candidates. The table is estimated as follows.

1. For each raw file, MS2 scans are selected at random and queried against the fragment index in an accumulator mode that, rather than emitting candidates, tallies for every pattern the number of times it arises from target and from decoy precursors. For speed, patterns matching fewer than two fragments are excluded from consideration.
2. For each pattern, Pioneer computes a library-corrected excess rate—the relative excess of target over decoy counts. A pattern passes if the rate exceeds 3%.
3. If a file yields fewer than one million total counts, per-pattern rates are considered unreliable and all patterns are instead pooled into 18 coarse groups, defined by whether the top-ranked fragment was observed and by how many of the top three and top five were observed. All patterns in a group share the verdict computed from the group's pooled counts.

#### 1.5 Parameter Tuning

Pioneer performs a preliminary analysis of each raw data file to estimate run-specific parameters for library retention time alignment, mass accuracy, and library collision energy alignment. This "pre-search" follows a similar procedure to the main search but with three modifications. First, Pioneer samples a subset of MS2 scans until it has collected a target number of peptide-spectrum-matches (PSMs) at a 1% FDR threshold. The MS2 scans are partitioned into retention time bins, and the sampler cycles through the bins in turn, each time taking the highest total-ion-current scan remaining in the current bin. Second, the tuning search requires a precursor's top four ranked fragments to match in the fragment index, together with a minimum total number, initially eight, of matched fragments. Third, the tuning search employs a simplified version of the fragment index that contains only a single retention time bin.

The pre-search proceeds in two phases. A wide scout phase searches with a deliberately wide fixed mass tolerance (±100 ppm) to estimate the m/z bias and dispersion; a collection phase then re-searches with the fitted mass-error model to gather the PSMs used for the remaining models. In each phase the number of sampled scans is increased adaptively from the observed PSM rate; if the PSM target is not reached, the minimum total number of matched fragments is lowered and the search repeated over all scans for up to three successive reductions. The parameter tuning search estimates the following run-specific parameters:

- A uniform-basis cubic B-spline that maps empirical retention times to library retention times.
- A retention time tolerance measured in library retention time units.
- An MS2 mass-error model that expresses the m/z bias and tolerance as smoothing-spline functions of fragment m/z, intensity, and retention time.
- An MS1 m/z error and tolerance (ppm), estimated from the precursor isotope pattern of the accepted PSMs.
- A function that returns an optimized library collision energy given the m/z and charge of a precursor.

##### 1.5.1 Retention Time Alignment

Let $\mathcal{D} = \{(t_i^{\text{emp}}, t_i^{\text{lib}})\}_{i=1}^{n}$ be the set of matched PSMs in the pre-search that pass a 1% FDR threshold, retaining at most the three highest-scoring PSMs per precursor. For each PSM

- $t_i^{\text{emp}}$ is the empirical retention time
- $t_i^{\text{lib}}$ is the library retention time
- $n$ is the number of sampled PSMs

Pioneer fits a uniform-basis cubic B-spline $f(t)$ that maps empirical to library retention times by penalized least squares; outlier PSMs identified from an initial fit are removed before the spline is refit, and the refit spline is constrained to be monotonic. The retention time errors of the retained PSMs are

$$\epsilon_i = t_i^{\text{lib}} - f(t_i^{\text{emp}}) \tag{2}$$

Pioneer sets the retention time tolerance to 4 times a robust estimate of standard deviation. Pioneer calculates the median absolute deviation and retention time tolerance, $\delta_{\text{RT}}$, as follows:

$$\text{MAD} = \text{median}\left(|\epsilon_i - \text{median}(\epsilon)|\right) \tag{3}$$

and

$$\delta_{\text{RT}} = \frac{4}{\Phi^{-1}(3/4)} \cdot \text{MAD} \tag{4}$$

##### 1.5.2 Mass Error Estimation

For each PSM, Pioneer selects the singly-charged y-ions of ion position four or greater among the three highest-ranked matched library fragments. The ranking of library fragment abundances is determined by the area-under-the-curve of the Altimeter splines. For each such fragment, Pioneer records its theoretical m/z, observed m/z, and intensity, together with the retention time of the scan. Mass errors are modeled in daltons,

$$\text{Error}(f_j) = m_j^{\text{measured}} - m_j^{\text{theoretical}}. \tag{5}$$

Pioneer models the systematic component of the mass error—the bias—as an additive function of fragment m/z, log-intensity, and retention time:

$$\text{bias}(m, I, t) = g_m(m) + g_I(\log_2 I) + g_t(t), \tag{6}$$

where each term is a regularized cubic B-spline fit by robust (iteratively reweighted) regression to down-weight outliers. The terms are fit in sequence, each to the residuals left by the previous. The m/z and retention time terms are fit to the median error within equal-count bins of m/z and retention time, while the intensity term is fit to all matched fragments under a monotonicity constraint. The retention time term is fit only when at least 500 matched fragments span a non-degenerate retention time range, and is otherwise set to zero. Each bias spline is evaluated by linear extrapolation outside its fitted range. During the search, every predicted fragment m/z is shifted by the fitted bias before matching.

The mass tolerance is derived from the dispersion of the residual errors rather than from a single fixed window. Pioneer fits a smoothing spline giving a robust estimate of the residual standard deviation as a function of log-intensity, together with an m/z-dependent multiplicative correction, and sets the half-width of the matching window to $k$ standard deviations, with $k$ fit per run to cover approximately 95% of the residual errors. Both spread splines are held constant outside their fitted range.

When too few fragments are matched to fit these splines, Pioneer falls back to a simpler model with a constant bias and fixed left- and right-hand tolerances in ppm.

##### 1.5.3 MS1 Mass Error Estimation

Pioneer also estimates an MS1 mass-error model from the same MS2-accepted PSMs. For each PSM, Pioneer locates the MS1 scan nearest in retention time and searches it for the precursor's first three isotopologues (M+0, M+1, M+2). For each isotopologue, the closest MS1 peak within ±100 ppm of the predicted m/z is matched, and the signed ppm residual between the observed and predicted m/z is recorded. The MS1 mass bias is set to the median of these residuals and the MS1 tolerance to ±5 times a robust estimate of their standard deviation, $\text{MAD}/\Phi^{-1}(3/4)$. Unlike the MS2 model, the MS1 model is a single bias and tolerance rather than a set of splines. Pioneer uses it to correct and bound precursor m/z in the MS1-level steps of the search.

##### 1.5.4 Library Collision Energy Alignment

Pioneer selects an optimized collision energy as a function of precursor m/z and charge that best aligns the Altimeter spectral library to the data for each run in an experiment. Recall that the Altimeter libraries include fragment intensity splines for each precursor that can be evaluated at different normalized collision energies (NCEs). During the search, Pioneer evaluates these splines at specific NCE values to get fragment intensities for the search. For each precursor in a set of pre-search PSMs, Pioneer evaluates the library at every value on a grid of 20 NCEs spanning 21 to 40 and scores the resulting fit to the spectrum by the deconvolution goodness of fit. The NCE giving the best fit is taken as that precursor's optimal value:

$$\text{NCE}_p = \underset{n \in \mathcal{N}}{\text{argmax}}\ \text{gof}(p, n). \tag{7}$$

Pioneer then summarizes these per-precursor optima into a model indexed by charge state and precursor m/z. For each charge state, the optimal NCE values are binned by precursor m/z and the median NCE is taken within each bin; the number of m/z bins is reduced adaptively until every bin contains at least 50 precursors. The model predicts a precursor's NCE as the median of the bin matching its charge and m/z. Charge states with fewer than 50 precursors are not fit; a precursor of such a charge is assigned the value from the nearest charge state that was fit, or a default NCE if no charge state was. Pioneer uses this model to assign NCE values for all precursors in subsequent searches.

#### 1.6 Main Search

In the main search, Pioneer identifies, scores, and filters the precursors carried forward for quantification. The main search begins with its own fragment-index search (Section 1.4), retrieving candidate precursors for each MS2 scan. This is a separate search from the one performed during parameter tuning (Section 1.5): parameter tuning queries a simplified index with a single retention time bin and a wide mass tolerance, whereas the main search queries the full index—using all retention time bins together with the calibrated retention time alignment and mass-error model fit during tuning—to constrain candidates by both retention time and m/z. Each scan's candidate precursors are then quantified against the spectrum by spectral deconvolution (Section 1.7) and scored by a LightGBM classifier. Pioneer then selects a single best PSM per precursor, and removes low-confidence identifications.

##### 1.6.1 PSM Scoring

For each MS data file, Pioneer scores its PSMs with a gradient-boosted decision-tree classifier (LightGBM) trained to separate target from decoy PSMs. Training uses two-fold cross-validation: precursors are split into two folds, a model is trained on each fold and used to score the PSMs of the other, so no PSM is scored by a model trained on it. The classifier draws on a large set of features, including spectral goodness-of-fit and similarity scores, fragment-ion coverage, the deconvolution weight and its rank among co-eluting precursors, retention-time agreement, MS1 isotope-pattern features, per-rank fragment chromatographic traces and fragment–fragment correlations, and precursor and sequence properties. For files with too few PSMs to train a stable tree model, Pioneer also trains a probit regression and keeps whichever classifier identifies more targets at a 1% out-of-fold FDR. From these scores, Pioneer reduces each precursor to a single best PSM and discards those with a posterior error probability above 0.99.

##### 1.6.2 Retention Time Refinement and Tolerance Estimation

After per-file scoring and PEP filtering, each run's surviving precursors are split by cross-validation fold and carried forward for re-scoring, but first, Pioneer uses the main search results to refine its retention time models in three ways.

***Refinement of predicted retention times.***

First and for each file, Pioneer finds the best scoring PSM per precursor. From the best PSMs, Pioneer collects the target precursors identified at a per-file, q-value $q \leq 0.015$, and regresses their retention time residuals—observed iRT minus predicted iRT—by ordinary least squares on the predicted iRT, its square, and counts of tokens encoding each residue together with its modification state, supplemented by tokens identifying the peptide's N- and C-terminal residues. A model is fit for each cross-validation fold using only precursors from the other fold, and is applied to the predicted retention times of every candidate PSM in its own fold. Pioneer then updates the retention-time-based features for each PSM, re-scores all candidate PSMs using the classifier already trained in the first pass, and then again finds the best PSM per precursor. The step is skipped when fewer than 250 high-confidence target precursors are available to fit a fold's model.

***Refinement of the empirical alignment.***

Second, Pioneer then replaces the retention time alignment fit during parameter tuning. For each run, the best PSM of every target precursor with a classifier score greater than 0.9 is used to refit the spline that maps empirical retention times to the refined library retention times (Section 1.5.1). The per-run tolerance, in library retention time units, is set to four times a robust estimate of the standard deviation of the residuals about the refit spline, $\mathrm{MAD}/\Phi^{-1}(3/4)$. Runs with fewer than 30 qualifying PSMs retain the model from parameter tuning.

***Retention time tolerance for chromatogram extraction.***

Finally, Pioneer estimates the retention time tolerance, shared by all runs, that bounds the window over which chromatograms are later extracted. For each precursor it computes the log-odds average of the top-$k$ of its cross-validated run probabilities $\{s_r(p)\}$,

$$s_{\text{global}}(p) = \sigma\left(\frac{1}{k}\sum_{r\in\text{top-}k}\log\frac{s_r(p)}{1-s_r(p)}\right), \qquad k = \min\Big(|\{s_r(p)\}|\,,\ \lfloor\sqrt{R}\rfloor\Big)\,, \tag{8}$$

where $R$ is the number of runs, $\sigma$ is the logistic function, and probabilities are clamped below at 0.02. These scores are used only here. Target precursors passing a target-decoy q-value of 0.001 are partitioned into 20 equal-occupancy bins of observed retention time, and within each bin the 100 precursors of largest deconvolution weight set the half-width

$$\delta_{\text{RT}} = \max(2\,\text{median}(\text{FWHM}),\ 10\,\tau)\,, \tag{9}$$

with $\tau$ the 75th percentile of their cycle times, each estimated as a precursor's FWHM divided by one less than its number of scans above half maximum.

#### 1.7 Matrix Representation of Library and Observed Mass Spectra

For each MS2 spectrum, Pioneer aligns it to candidate library spectra for precursors within a specified mass-to-charge (m/z) and retention time tolerance. Let $N$ denote the number of candidate precursors considered for the scan.

Let the observed spectrum contain $m$ peaks and be represented as

$$X = \{(I_k^{(X)}, Z_k^{(X)})\}_{k=1}^{m},$$

where $I_k^{(X)}$ and $Z_k^{(X)}$ denote the intensity and m/z of the $k$th observed peak.

For $j \in \{1, \ldots, N\}$, let the candidate library spectrum for precursor $j$ contain $n_j$ fragments and be represented as

$$L_j = \{(I_k^{(L_j)}, Z_k^{(L_j)})\}_{k=1}^{n_j},$$

where $I_k^{(L_j)}$ and $Z_k^{(L_j)}$ denote the intensity and m/z of the $k$th fragment in library spectrum $j$. All spectra are sorted in ascending order by m/z.

Pioneer combines all candidate library spectra into a single set

$$F = \{(I_k^{(F)}, Z_k^{(F)}, j_k^{(F)})\}_{k=1}^{T},$$

where $T = \sum_{j=1}^{N} n_j$, and $j_k^{(F)}$ identifies the precursor associated with each fragment.

After sorting $F$ by $Z^{(F)}$, Pioneer matches each fragment in $F$ to the nearest peak in $X$ based on m/z, provided the peak is within the specified mass tolerance. The matched and unmatched fragments are represented as:

$$F_m = \{(I_k^{(F_m)}, j_k^{(F_m)}, m_k^{(F_m)})\}_{k=1}^{K}$$

and

$$F_u = \{(I_k^{(F_u)}, j_k^{(F_u)})\}_{k=1}^{U},$$

where $K = |F_m|$ and $U = |F_u|$ denote the number of matched and unmatched library fragments, respectively. Here, $m_k^{(F_m)} \in \{1, \ldots, m\}$ denotes the index of the observed peak in $X$ matched to the $k$th library fragment.

Let

$$P = \left|\{m_k^{(F_m)}\}\right|$$

denote the number of unique peaks in $X$ matched by fragments in $F_m$. Let $U = |F_u|$ denote the number of unmatched fragments. The matrix row dimension is then

$$M = P + U,$$

where $P \leq K$.

Define a mapping $r : \{1, \ldots, K\} \to \{1, \ldots, P\}$ such that if two fragments $k$ and $l$ match to the same peak in $X$ (i.e., $m_k^{(F_m)} = m_l^{(F_m)}$), then $r(k) = r(l)$. In other words, matched fragments that correspond to the same observed peak share a common row in the matrix, whereas each unmatched fragment in $F_u$ is assigned a unique row.

The observed intensities vector $\vec{y} \in \mathbb{R}^M$ and the template matrix $\mathbf{A} \in \mathbb{R}^{M \times N}$ are constructed as:

$$y_i = \begin{cases} I^{(X)}_{m_k^{(F_m)}} & \text{if } \exists k \leq M_m \text{ such that } r(k) = i \\ 0 & \text{otherwise} \end{cases} \tag{10}$$

$$A_{ij} = \begin{cases} \sum_{k\,:\,r(k)=i,\; j_k^{(F_m)}=j} I_k^{(F_m)} & \text{if } i \leq P \\ I_{i-P}^{(F_u)} & \text{if } i > P \text{ and } j_{i-P}^{(F_u)} = j \\ 0 & \text{otherwise} \end{cases} \tag{11}$$

This construction can be visualized schematically as shown below, with the top block representing matched observed peaks and the bottom block representing unmatched library fragments that occupy their own rows.

$$\underbrace{\begin{bmatrix} y_1 \\ \vdots \\ y_P \\ \hline y_{P+1} \\ \vdots \\ y_{P+U} \end{bmatrix}}_{\vec{y}\in\mathbb{R}^M,\ M=P+U} = \underbrace{\begin{bmatrix} A_{11} & \cdots & A_{1N} \\ \vdots & \ddots & \vdots \\ A_{P1} & \cdots & A_{PN} \\ \hline A_{P+1,1} & \cdots & A_{P+1,N} \\ \vdots & \ddots & \vdots \\ A_{P+U,1} & \cdots & A_{P+U,N} \end{bmatrix}}_{\mathbf{A}\in\mathbb{R}^{M\times N}} \underbrace{\begin{bmatrix} x_1 \\ \vdots \\ x_N \end{bmatrix}}_{\vec{x}\in\mathbb{R}^N}. \tag{12}$$

Pioneer represents $\mathbf{A}$ in a sparse, column-major layout.

#### 1.8 Linear Regression of Mass Spectra onto Library Spectra

Pioneer models each spectrum as a linear combination of $N$ template spectra from a spectral library. This problem can be formulated as the following linear system:

$$\mathbf{A}\vec{x} \approx \vec{y} \tag{13}$$

Pioneer estimates a vector of weights, $\vec{x} \in \mathbb{R}^N$, subject to a non-negativity constraint. Because $\mathbf{A}$ is sparse, Pioneer uses a coordinate descent algorithm, which iteratively updates each variable $x_j$ to optimize the objective $L(\vec{x})$ while treating the other variables as constants. For each $j$, Pioneer solves this optimization problem:

$$x_j^{\mathrm{k}+1} = \operatorname*{argmin}_{x\geq 0} L(x; x_1^{k+1}, \ldots, x_{j-1}^{k+1}, x_{j+1}^{k}, \ldots, x_N^{k}) \tag{14}$$

Pioneer enforces the non-negativity constraint by setting the new guess for each variable to the maximum of 0 and the updated value, i.e., $x_j = \max(0, x_j^{\mathrm{new}})$. Pioneer also uses a “hot-start”. That is, if a previous spectrum already estimated a weight for a given precursor, then that previous estimate is the initial guess.

Pioneer uses two objective functions at different stages of the analysis. During identification—the main search and quadrupole transmission tuning—it maximizes a Poisson log-likelihood. During chromatogram integration, where the weights are the quantitative readout, it minimizes a pseudo-Huber loss. The parameter tuning pre-search, which requires only approximate weights, minimizes the sum of squared residuals.

##### 1.8.1 Poisson Objective

Writing $\vec{\mu} = \mathbf{A}\vec{x}$ for the predicted peak intensities, Pioneer maximizes the log-likelihood of an identity-link Poisson model without intercept:

$$\operatorname*{argmax}_{\vec{x}\geq \mathbf{0}} \ell(\vec{x}) = \sum_{i=1}^{M} \left[y_i \log \mu_i - \mu_i\right] \tag{15}$$

Each coordinate is updated by a single Newton step, computed from the first derivative of the negative log-likelihood along coordinate $j$ and the corresponding observed information,

$$L1_j = \sum_{i=1}^{M} A_{ij}\left(1 - \frac{y_i}{\mu_i}\right), \qquad L2_j = \sum_{i=1}^{M} A_{ij}^2 \frac{y_i}{\mu_i^2}, \tag{16}$$

$$x_j \leftarrow \max\left(x_j - \frac{L1_j}{L2_j},\ 0\right). \tag{17}$$

Because $\mathbf{A}$ is stored column-sparse, both sums run only over the peaks that precursor $j$ explains, and $\vec{\mu}$ is updated incrementally after each step rather than recomputed. When a column’s observed information vanishes—which happens when every peak it explains has zero intensity—Pioneer substitutes the expected (Fisher) information, $\sum_i A_{ij}^2/\mu_i$, so that the step remains defined.

##### 1.8.2 Pseudo-Huber Objective

Let

$$\vec{r} = \mathbf{A}\vec{x} - \vec{y} \tag{18}$$

be the vector of residuals, $\vec{r} \in \mathbb{R}^M$. Pioneer minimizes the pseudo-Huber loss [7, 8]:

$$\underset{\vec{x}\geq \mathbf{0}}{\operatorname{argmin}}\, L(\vec{x}) = \delta^2 \left( \sum_{i=1}^{M} \sqrt{1 + \left(\frac{r_i}{\delta}\right)^2} \right) - \delta^2 M \tag{19}$$

The smoothing parameter $\delta$ controls the transition between squared and absolute error. Pioneer solves each single variable problem by the Newton-Raphson method. In case Newton's method fails to converge, Pioneer defaults to a bisection method, bracketing the root between 0 and the largest iterate at which the derivative remained positive, capped at a large positive value ($10^{11}$).

***Calibration of $\delta$***

Pioneer estimates a single $\delta$ for the experiment before integration. For a sample of up to 5,000 PSMs, each spectrum is re-solved over a geometric grid of candidate $\delta$ values, and each precursor's weight is recorded as a function of $\delta$. Curves whose weight varies by less than 10% across the grid are discarded as uninformative. For the remainder, Pioneer takes the $\delta$ at which the weight has completed half of its total change, and the median of these values across all curves becomes the experiment's $\delta$.

#### 1.9 Precursor Scoring, Match-Between-Runs, and False-Discovery Rate Control

Pioneer re-scores the best PSM per precursor from each file from the main search. Scoring proceeds in three stages that use gradient-boosted decision tree models (LightGBM) to discriminate targets from decoys. When match-between-runs (MBR) is enabled, precursors that are well supported globally but score poorly in a given run are re-examined after chromatogram integration. If those precursors survive a counterfactual test they are recovered under a constraint on the combined error rate.

##### 1.9.1 Data Structure and Cross-Validation Setup

Pioneer re-scores the best-scoring PSM for each precursor, $p$, in run, $r$, as described by a feature vector $\mathbf{X}_{p,r} \in \mathbb{R}^d$, which summarizes that precursor's spectral match, chromatographic peak, and the behavior of the co-eluting precursors it competes with. Each precursor carries a label $Y_p \in \{0, 1\}$ that indicates its target (1) or decoy (0) status.

During spectral library construction, Pioneer assigns each library protein group at random to one of two cross-validation folds, $\mathcal{F}_1$ or $\mathcal{F}_2$. Every precursor inherits the fold of its protein group.

##### 1.9.2 Semi-Supervised Scoring of Precursor-Run Observations

Pioneer trains a LightGBM classifier to separate target from decoy observations using spectral, chromatographic, and retention time features. Training is semi-supervised and iterative.

Let $s_{p,r}^{(t)}$ denote the cross-validated (out-of-fold) probability at iteration $t$, and $q_{p,r}^{(t)}$ the corresponding target-decoy q-value. The training set at iteration $t$ is

$$\mathcal{T}^{(t)} = \{(p, r) : Y_p = 0\} \,\cup\, \left\{(p, r) : Y_p = 1,\; q_{p,r}^{(t-1)} \leq 0.03\right\}, \tag{20}$$

that is, all decoy observations, and then all target observations with better than a 3% q-value at the previous iteration. Two models are fit per iteration, one per cross-validation fold, and each scores the observations of the held-out fold.

Iteration continues until the number of target observations passing a 1% q-value fails to improve by at least 1% over the previous iteration, or until a maximum of eight iterations. The iteration yielding the most target observations at 1% q-value is retained, and its out-of-fold probabilities become the per-run precursor probabilities $\text{prec\_prob}_{p,r}$ used downstream. When MBR is enabled, Pioneer additionally retains the in-fold probabilities, which serve as a feature for the transfer model described below.

##### 1.9.3 Identification-based run similarity

After per-run precursor scoring, Pioneer computes a cross-run similarity metric based on precursor identifications. Let $I_r$ be the set of precursors indices passing the experiment-wide q-value threshold. For precursor $i$, let $f_i$ be the number of runs containing $i$, and assign the inverse-frequency weight

$$w_i = \log\left(\frac{|R|+1}{f_i+1}\right). \tag{21}$$

The directional similarity from receiver run $r$ to donor run $d$ is

$$\mathrm{sim}(r \to d) = \frac{\sum_{i \in I_r \cap I_d} w_i}{\sum_{i \in I_r} w_i}. \tag{22}$$

The similarity metric measures how completely the weighted precursor set observed in the receiver is represented in the donor. The run similarities are used in global scoring and match-between-runs.

##### 1.9.4 Global Precursor Score

Pioneer estimates error rates in the experiment-wide and global contexts of Rosenberger et al. [9]. Pioneer calculates a global score by training a global model on a set of cross-run summary statistics. The summary features comprise the largest three per-run LGBM scores for each precursor and the gaps between them; their mean, median, standard deviation, and minimum; the number of runs in which the precursor was identified and the number exceeding probability thresholds of 0.5, 0.9, and 0.99; three measures of how the observed runs sit within the experiment's run-similarity structure; and log-odds combinations of the top two and top three probabilities. The log-odds combination of the top $k$ probabilities is

$$\mathrm{logodds}_k(p) = \sigma\left(\frac{1}{k}\sum_{r \in \text{top-}k} \log\frac{\mathrm{prec\_prob}_{p,r}}{1-\mathrm{prec\_prob}_{p,r}}\right), \tag{23}$$

where $\sigma$ is the logistic function and probabilities are clamped to $[0.1, 1-10^{-6}]$ before the transformation.

Pioneer trains a LightGBM model out-of-fold on these features to globally discriminate target from decoy precursors, constraining the model to be monotone in the features for which additional evidence must not lower the score. As a safeguard, Pioneer also forms a purely empirical score—the log-odds combination over the top $\lfloor\sqrt{|\mathcal{R}|}\rfloor$ runs, where $\mathcal{R}$ is the set of runs in the experiment—which is supplied to the model as a feature and also retained as a fallback. Pioneer keeps the learned score only if it yields more target identifications at the target q-value threshold, counting each passing precursor; ties favor the empirical score. The retained value is the global precursor score $\mathrm{global\_prob}_p$.

##### 1.9.5 Q-values and Initial Filtering

Pioneer controls the false discovery rate at two levels, global and experiment-wide. Global q-values are computed by target-decoy competition over $\mathrm{global\_prob}_p$; experiment-wide q-values are computed over $\mathrm{prec\_prob}_{p,r}$, ranking each precursor-run pair. A precursor-run pair enters the confident set only if it satisfies both thresholds,

$$\mathcal{P}_{\text{pass}} = \left\{(p,r) : \mathrm{global\_qval}_p \le \alpha_g \text{ and } \mathrm{qval}_{p,r} \le \alpha_e\right\}, \tag{24}$$

with $\alpha_g$ and $\alpha_e$ both set to 0.01 by default. When MBR is disabled, the experiment-wide q-values are then recomputed over the passing subset alone, and the analysis proceeds to quantification. When MBR is enabled, recomputation is deferred until after transfers have been accepted.

#### 1.10 Chromatogram Smoothing and Integration

For target precursors passing both global and experiment-wide q-value thresholds, Pioneer estimates their chromatographic peak area. Pioneer does this by first repeating the linear regression of mass spectra onto the library spectra (Section 1.8), but with the now substantially reduced list of identified precursors. The precursor-specific weights are sorted in temporal order to form chromatograms, which are processed and integrated as follows.

##### 1.10.1 Isotope Trace Handling

Precursor isotopic envelopes may be split between multiple quadrupole isolation windows in a DIA method. The result is multiple chromatograms per precursor, which we refer to as isotope traces. The `combine_traces` setting governs how isotope traces are incorporated into precursor chromatograms for quantification. The two strategies are:

***Combined Isotope Traces (combine_traces: true)***

When `combine_traces` is enabled, Pioneer combines intensities across all isotope traces for each precursor into a single chromatogram before integration. See section 1.13 for documentation on correcting the abundance of separate isotope traces based on the isolated precursor fraction.

***Separate Isotope Traces (combine_traces: false)***

When `combine_traces` is disabled, Pioneer instead quantifies from a single isotope trace per precursor. It selects the trace with the highest transmitted precursor fraction, breaking ties deterministically, and integrates only that trace, discarding the points belonging to the others.

##### 1.10.2 Integration Workflow

For each target precursor passing FDR thresholds, Pioneer applies the following chromatogram processing pipeline:

1. **Whittaker-Henderson Smoothing**: Apply penalized least squares smoothing [10] to the intensity time series. Each point is first divided by the fraction of the precursor's isotopic envelope transmitted by the isolation window that produced it. That fraction is the weight in the penalized fit. If the fraction is below 0.25 the weight is set to zero.
2. **Apex Refinement**: Start at the best PSM identified in the main search, and then climb the smoothed chromatogram to the local maximum in each direction. Treat whichever local maximum is higher as the apex.
3. **Peak Boundary Detection**: Identify the peak integration boundaries. Start two chromatogram points before and after the apex, and advances to the first local maximum of the second discrete derivative of the smoothed signal. Continue outward so long as the intensity does not exceed 15% above the running minimum. The boundary is placed at the running minimum after stopping.
4. **Baseline Subtraction**: Subtract a linear baseline drawn between the smoothed intensities at the two integration boundaries and clamp any negative values to zero.
5. **Trapezoidal Integration**: Integrate the baseline-corrected, smoothed time series by the trapezoidal rule, with a half-triangle contributed by each endpoint:

$$A_{p,r} = \frac{1}{2}\left[\tau(I_1 + I_N) + \sum_{j=1}^{N-1}(t_{j+1} - t_j)(I_j + I_{j+1})\right] \tag{25}$$

   where $A_{p,r}$ is the chromatographic peak area of precursor $p$ in run $r$, $t_j$ are retention times, $I_j$ are baseline-corrected intensities, $\tau$ mean retention time spacing between chromatogram points, and $N$ is the number of points within the integration bounds.
6. **Quantifiability Filter**: Discard noisy peaks. If the smoothed apex is not positive after baseline subtraction, or if the area over the same window before baseline subtraction is at least five times the area after baseline subtraction, Pioneer reports an area of zero.

The integrated peak areas $A_{p,r}$ serve as precursor abundances for downstream label-free quantification. A precursor may therefore pass false-discovery control and still carry no quantity in a given run; downstream quantification treats a zero area as an absent measurement rather than as a measurement of zero.

#### 1.11 Match-Between-Runs

After chromatogram integration, match-between-runs (MBR) may recover PSMs that passed the global q-value threshold but failed the experiment-wide threshold.

##### 1.11.1 Candidate and Donor Selection

Donors are PSMs that passed both the global and experiment-wide q-value thresholds. Candidates are PSMs that scored below the experiment-wide threshold but where the precursor has a donor PSM from a different run. For each candidate PSM, Pioneer identifies every run in which that precursor has an eligible donor. When multiple donor runs are available, Pioneer selects the donor from the run having the greatest directional similarity to the candidate's receiver run, as defined in Section 1.9.3; ties are broken by the donor's score.

Each candidate PSM is then evaluated four times: once against its true donor and again against three counterfactual donors. A counterfactual donor is the donor PSM of a different precursor, drawn from the same run as the true donor, among precursors of identical charge and length, and nearest to the candidate in predicted retention time. When that run cannot supply three counterfactual donors, Pioneer draws on progressively coarser pools drawn from the whole experiment.

##### 1.11.2 Transfer Features

Pioneer trains a semi-supervised match-between-runs model to recover the candidate PSMs. The model uses three kinds of features: receiver features, paired features, and cluster features.

Receiver features describe the candidate PSM alone and are carried forward from the main search and precursor scoring. These include the PSM's within-fold LightGBM score, spectral goodness-of-fit and distance measures, retention time error, deconvolution weight, the weight rank among co-eluting precursors, the fraction of the precursor isotope envelope transmitted, and sequence length.

Paired features compare the candidate PSM to the donor PSM. Most of these features are computed during chromatogram integration rather than the main search. The features include the log ratio of the integrated intensities, the log ratio of the explained spectral intensities, the difference between retention time residuals from prediction, the difference between observed apex retention times, the difference in numbers of integrated points, and the agreement between fragment intensity patterns. Two additional features are the donor's score from the semi-supervised scoring model (Section 1.9.2), and the similarity between the candidate's and donor's runs.

Cluster features place the candidate's run among the runs most similar to it. Runs are clustered on their intensity profiles, and Pioneer records how many of the other runs in the candidate's cluster identified the precursor, how many such runs there are, and the ratio of the two.

##### 1.11.3 The Transfer Model

Pioneer trains a LightGBM model that follows the same semi-supervised scheme as the experiment-wide scoring model (Section 1.9). The positive class is all target candidates on the first iteration, and then on the second iteration, all target candidates scoring better than a 3% q-value in the prior iteration; the negative class is candidates of decoy PSMs together with every candidate's three counterfactual rows. Iteration stops when the number of accepted target transfers fails to improve by 1%, and the best of at most eight iterations is retained. Each candidate then receives a score from its true-donor row, $s^{\text{true}}$, and another score that is the maximum from its three counterfactual rows, $s^{\text{false}}$.

##### 1.11.4 False Transfer Rate Control

A target candidate is counted as a false-transfer event when both $s^{\text{true}}$ and $s^{\text{false}}$ exceed the acceptance threshold, and the event is assigned the score $\min(s^{\text{true}}, s^{\text{false}})$. Let $T_0$ and $D_0$ be the numbers of target and decoy precursor-run pairs already passing without MBR. For a score threshold $\tau$, let $T(\tau)$ and $D(\tau)$ be the numbers of target and decoy candidates recovered at that threshold and $F(\tau)$ the number of false-transfer events above it:

$$\text{FTR}(\tau) = \frac{D_0 + D(\tau) + F(\tau)}{T_0 + T(\tau)}. \tag{26}$$

Pioneer computes this rate over the sorted candidate scores, enforces monotonicity from the lowest score upward to obtain a q-value for each candidate, and recovers those with a q-value at or below $\alpha$. No transfers are recovered if the baseline error rate $D_0/T_0$ already exceeds $\alpha$. Recovered candidates are merged back into the passing set, and the experiment-wide q-values and posterior error probabilities are refit over the combined set.

#### 1.12 Protein Inference and Quantification

After precursor-level FDR control, Pioneer performs protein inference globally. The inferred protein groups may be distinct from the library protein groups (see Section 1.3) and are hereafter simply referred to as protein groups.

##### 1.12.1 Input Selection for Protein Inference

Pioneer performs protein inference on the union, across all runs, of the precursors retained after the two-stage FDR control described in Section 1.9.5 and after match-between-runs (Section 1.11). Let $\mathcal{P}_{\text{pass}}$ denote the set of retained precursor-run pairs and $\mathcal{P}_{\text{inference}}$ the set of precursors eligible for protein inference:

$$\mathcal{P}_{\text{inference}} = \{p : \exists\, r \in \mathcal{R} : (p, r) \in \mathcal{P}_{\text{pass}}\} \tag{27}$$

Both target precursors ($Y_p = 1$) and decoy precursors ($Y_p = 0$) are included. Precursors are reduced to unique peptide sequences before inference, so that different charge states of the same sequence are treated as a single peptide.

##### 1.12.2 Parsimony-Based Protein Inference Algorithm

Pioneer implements a two-phase parsimony-based inference algorithm [11, 12] to identify a minimal set of protein groups that can explain all the observed peptides. The algorithm operates on the peptide–protein bipartite graph defined below. Peptides are first partitioned by target/decoy status and entrapment group, and the algorithm is applied independently within each of these populations, so that a protein group never contains peptides of mixed status.

***Protein-Peptide Bipartite Graph***

Let $\mathcal{A}$ denote the set of all protein accession numbers in the spectral library (Section 1.3). Define the set of peptide-protein associations:

$$\mathcal{E} = \{(p, a) : p \in \mathcal{P}_{\text{inference}}, a \in \mathcal{A}, \text{peptide } p \text{ maps to protein } a\} \tag{28}$$

Each pair $(p, a) \in \mathcal{E}$ represents an edge in a bipartite graph connecting observed peptide $p$ to library protein name $a$.

***Algorithm Overview***

The inference algorithm decomposes the bipartite graph into disjoint connected components using depth-first search. Within each component, the algorithm applies a two-phase approach. Pioneer first selects all proteins with unique peptide evidence and next applies a greedy set-cover. At each iteration of the greedy phase, inferred proteins with identical remaining peptide sets are first merged, and the inferred protein group covering the most remaining peptides is selected. The algorithm returns only peptides that uniquely map to a single protein group; shared peptides are excluded to ensure unambiguous protein quantification.

**Algorithm 1** Protein Inference Phase 1: Graph Decomposition

**Input:** Edges $\mathcal{E} = (p, a)$ where $p$ is peptide, $a$ is protein
**Output:** Connected components $C = (P_c, A_c)$ with bidirectional mappings $A[p]$ and $P[a]$
**Define:** $A[p] = a : (p, a) \in \mathcal{E}$ (proteins containing peptide $p$)
    $P[a] = p : (p, a) \in \mathcal{E}$ (peptides in protein $a$)

*// Build bidirectional mappings*
**for** $(p, a) \in \mathcal{E}$ **do**
    $A[p] \leftarrow A[p] \cup a$ ▷ Proteins for each peptide
    $P[a] \leftarrow P[a] \cup p$ ▷ Peptides for each protein
**end for**

*// Find connected components via depth-first search*
$V \leftarrow \emptyset$, $C \leftarrow \emptyset$ ▷ Visited peptides, components list
**for** each peptide $p$ **do**
    **if** $p \notin V$ **then**
        $(P_c, A_c) \leftarrow \text{DFS}(p, A, P, V)$ ▷ Discover component
        $V \leftarrow V \cup P_c$ ▷ Mark component peptides as visited
        $C \leftarrow C \cup (P_c, A_c)$
    **end if**
**end for**

**return** $C$, $A$, $P$

**Algorithm 2** Protein Inference Phase 2: Component Processing with Merge-First Set Cover

**Input:** Components $C$, mappings $A[p]$ and $P[a]$ from Algorithm 1
**Output:** $G[p] \to q$ where $q \subseteq \mathcal{A}$ is a protein group (only unique peptides)

**function** MERGEINDISTINGUISHABLE($K$, $P$, $R$)
    **Input:** Candidate proteins/groups $K$, peptide mapping $P[\cdot]$, remaining peptides $R$
    **Output:** Updated $K$ and $P[\cdot]$ with indistinguishable candidates merged
    Group candidates $a \in K$ by the set $P[a] \cap R$
    **for** each group $\Gamma$ with $|\Gamma| > 1$ **do**
        Create merged group identifier $g \leftarrow$ MergeName($\Gamma$) ▷ e.g., "protein1;protein2;..."
        $P[g] \leftarrow \bigcup_{a \in \Gamma} P[a]$ ▷ Define peptide set for merged group
        $K \leftarrow (K \setminus \Gamma) \cup \{g\}$
    **end for**
    **return** $K$, $P$
**end function**

*// Process each component independently*
**for** $(P_c, A_c) \in C$ **do**

    *// Case 1: All proteins indistinguishable*
    **if** $P[a] = P[a']$ for all $a, a' \in A_c$ **then**
        **for** $p \in P_c$ **do**
            $G[p] \leftarrow A_c$ ▷ All peptides unique to protein group
        **end for**
        **continue**
    **end if**

    *// Case 2: Mixed peptide assignments*
    $U \leftarrow p \in P_c : |A[p] \cap A_c| = 1$ ▷ Unique peptides

    *// Phase 1: Select all proteins with unique peptides*
    $S \leftarrow a \in A_c : P[a] \cap U \neq \emptyset$ ▷ Necessary proteins
    $R \leftarrow P_c \setminus \bigcup_{a \in S} P[a]$ ▷ Remaining uncovered peptides

    *// Phase 2: Merge-first greedy set cover*
    $K \leftarrow A_c \setminus S$ ▷ Candidate proteins
    **while** $R \neq \emptyset$ and $K \neq \emptyset$ **do**
        $K \leftarrow$ MergeIndistinguishable($K, P, R$) ▷ Merge proteins with identical remaining peptides
        $a^* \leftarrow \arg\max_{a \in K} |P[a] \cap R|$ ▷ Select protein covering most peptides
        **if** $|P[a^*] \cap R| = 0$ **then**
            **break**
        **end if**
        $S \leftarrow S \cup a^*$ ▷ Add to solution
        $R \leftarrow R \setminus P[a^*]$ ▷ Remove covered peptides
        $K \leftarrow K \setminus a^*$ ▷ Remove from candidates
    **end while**

    *// Add only unique peptides to result*
    **for** $p \in P_c$ **do**
        $S_p \leftarrow A[p] \cap S$ ▷ Necessary proteins for this peptide
        **if** $|S_p| = 1$ **then**
            $G[p] \leftarrow S_p$ ▷ Unique assignment
        **end if**
        *// Note: Shared peptides ($|S_p| > 1$) are excluded from $G$*
    **end for**
**end for**

**return** $G$

After protein inference, the protein groups are filtered by the minimum number of peptides. For each protein $q \in S$ selected across all components, let $\mathcal{N}(q)$ denote its set of assigned peptides. Filter proteins by:

$$S_{\text{filtered}} = q \in S : |\mathcal{N}(q)| \geq n_{\text{min}} \tag{29}$$

where $n_{\text{min}}$ is the minimum peptide threshold (default: 1).

##### 1.12.3 Protein Group Scoring and FDR Control

After protein inference, Pioneer scores protein groups in three stages that mirror the precursor-level procedure: a summary score computed from the constituent peptides, a learned per-run refinement, and a learned experiment-wide score.

###### *Stage 1: Per-Run Peptide Roll-Up*

In each run, Pioneer scores a protein group by summing $-\log(1 - p + \epsilon)$ over the precursors whose peptides were uniquely assigned to that group,

$$\text{pg\_score}(q) = -\sum_{p \in \mathcal{N}(q)} \log\left(1 - \text{prob}_p + \epsilon\right) \tag{30}$$

where $\mathcal{N}(q)$ is the set of contributing precursors, $\text{prob}_p$ is the precursor's probability, and $\epsilon$ is a small pseudocount that bounds the contribution of any single precursor. Only precursors flagged as usable for protein quantification and still passing both the run-specific and global precursor q-value thresholds contribute.

###### *Stage 2: Per-Run Model Refinement*

Pioneer refines the per-run score with a gradient-boosted decision tree model (LightGBM) trained to discriminate target from decoy protein groups. In addition to the roll-up score, the model uses the score obtainable from peptides that could not be assigned unambiguously, several measures of the fraction of a protein group's possible peptides that were observed, a measure of the consistency of the constituent precursors' chromatographic shapes, and indicators of whether the group rests on a single non-transferred peptide or only on transferred ones.

Training is semi-supervised and cross-validated on the folds inherited from precursor scoring. The roll-up score used directly when a dataset provides too few protein groups or too few decoys to fit a stable model.

###### *Stage 3: Global Protein Group Scoring*

Pioneer then computes a single score per protein group for the whole experiment, following the same construction as the global precursor score (Section 1.9.4). Cross-run summary statistics of the refined per-run scores—their three largest values and the gaps between them, log-odds combinations of the top two and top three, their mean, median, standard deviation and minimum, the number of runs in which the group was identified and the number exceeding fixed score thresholds, peptide counts and coverage summarized across runs, and measures of where the observed runs sit within the experiment's run-similarity structure—are combined by a LightGBM model. As with precursors, Pioneer also calculates a log-odds score over the top $\lfloor\sqrt{|\mathcal{R}|}\rfloor$ runs, supplies it to the model as a feature, and keeps the learned score only if it yields more target identifications at the target q-value threshold, counting each passing group once per run in which it passes.

###### *Protein-Level Q-value Calculation*

Pioneer controls both the experiment-wide and global false discovery rate at the protein group level. For a given score type (experiment-wide or global), let $\pi$ be the permutation sorting protein groups by decreasing score. The protein-level q-value for the $k$-th ranked protein is:

$$\text{protein\_qval}_{\pi(k)} = \min_{j \geq k} \left\{ \frac{\sum_{l=1}^{j}(1 - Y_{\pi(l)})}{\sum_{l=1}^{j} Y_{\pi(l)}} \right\} \tag{31}$$

where $Y_q = 1$ if protein $q$ is a target and $Y_q = 0$ if it is a decoy. After filtering on both thresholds the experiment-wide q-values are recomputed using the experiment-wide scores of the remaining protein groups and only these proteins are retained for downstream quantification.

##### 1.12.4 MaxLFQ Quantification

Pioneer implements the MaxLFQ algorithm [13] for label-free protein quantification. For each protein $q$ passing protein-level FDR thresholds and peptide $p \in \mathcal{N}(q)$ in run $r$, the integrated chromatographic peak area $A_{p,r}$ serves as the quantification intensity. The MaxLFQ intensities $\{I_{q,r}\}_{r\in\mathcal{R}}$ are computed by solving:

$$\min_{\{I_{q,r}\}_{r\in\mathcal{R}}} \sum_{r,r'\in\mathcal{R}} \sum_{p\in\mathcal{N}(q)} w_{p,r,r'} \left( \log I_{q,r} - \log I_{q,r'} - \log \frac{A_{p,r}}{A_{p,r'}} \right)^2 \tag{32}$$

where $w_{p,r,r'} = 1$ if $A_{p,r} > 0$ and $A_{p,r'} > 0$, and

#### 1.13 Fragment Isotope Correction

By default Pioneer assumes the spectral libraries provided to it record the total intensity rather than the monoisotopic intensity of each fragment ion. Pioneer therefore models the fragment isotope probabilities as conditional on the quadrupole-filtered precursor isotope distribution, which differs from the natural distribution. Using the conditional model, Pioneer re-isotopes the library spectra on the fly to more closely match the empirical MS2 spectra.

Goldfarb et al. described how fragment isotope distributions depend on precursor isotope selection and then derived a formula to calculate conditional fragment isotope probabilities in terms of unconditional probabilities [14]. Borrowing notation, suppose $P$ and $F$ are random variables for the isotopic state of a precursor and fragment ion respectively, and likewise, $p$ and $f$ are the specific isotopic states. Then for fragmentation of a specific subset of the precursor isotopes, $\mathbf{p}$, the conditional fragment isotope probabilities can be written as:

$$\Pr\big(F = f \mid \bigcup_{p\in\boldsymbol{p}} P = p\big) \tag{33}$$

Goldfarb et al. proposed a means to calculate these conditional fragment isotope probabilities in terms of the unconditional ones, $\Pr(F = f)$, and then proposed an efficient approximation for the unconditional probabilities using sulfur-specific splines.

Pioneer uses this method and extends it to model non-uniform quadrupole transmission efficiency. Pioneer models the transmission efficiency with which the quadrupole transmits each precursor isotope. The quadrupole transmission function is defined as the following:

$$Q(z^p; c, w) = \Pr\big(P = p \mid Q\big) \tag{34}$$

where $z^p$ is the mass-to-charge ratio of isotope $p$. The parameters $c$ and $w$ are the center and nominal width of the isolation window in Thomsons respectively.

This function describes the probability that the quadrupole transmits a specific precursor isotope through the entire length of the quadrupole. Assuming a reasonable estimate for $Q(z; c, w)$, and given equation 3.5 from [14] we have the following:

$$\Pr\big(F = f|Q\big) = \Pr\big(F = f\big) \cdot \left( \sum_{p=0}^{\infty} \Pr\big(P = p \mid Q\big) Pr\big(C = p - f\big) \right) \tag{35}$$

In practice, Pioneer truncates the sum in equation (1.13) at $p = 5$. Pioneer calculates fragment isotopes up to a user-defined maximum number of isotopes, which can be set differently for the first-pass search and the second-pass searches. Using this equation, Pioneer estimates conditional fragment isotope distributions assuming a reasonable model for $Q$.

#### 1.14 Estimating the Quadrupole Transmission Function

Pioneer estimates the quadrupole transmission function, $Q$, based on a random sampling of high-quality PSMs. Given a model for $Q$, it is possible to estimate the parameters based on deviations between observed and theoretical precursor isotope ratios. For precursor $i$, define:

- $\zeta_i^k$ - theoretical absolute abundance of the $k$-th precursor isotope prior to quadrupole isolation
- $x_i^k$ - observed abundance of the $k$-th precursor isotope after quadrupole isolation

- $z_i^k$ - m/z offset of the $k$-th precursor relative to the isolation window center
- $Q(z_i^k; c, w)$ - transmission probability at offset $z_i^k$ for a quadrupole centered at $c$, with an isolation width, $w$.

Then by definition:

$$x_i^k = \zeta_i^k \cdot Q(z_i^k; c, w) \tag{36}$$

For precursor $i$ define $\delta_i$:

$$\delta_i = \frac{\zeta_i^1}{\zeta_i^0} \tag{37}$$

Theoretical isotope ratios, $\zeta_i^1/\zeta_i^0$, can be calculated from the isotope splines provided in Goldfarb et al. That is, although mass spectrometers cannot measure the absolute number of each isotope in front of the quadrupole, it is possible to calculate the isotopic ratios from theory, either exactly based on the molecular composition or approximately via the averagine method or isotope splines [14]. Pioneer can then estimate the quadrupole transmission function, $Q(z_i^k; c, w)$, based on a comparison between the observed abundance ratios $x_k^0/x_k^1$ and the theoretical ratios $\zeta_i^0/\zeta_i^1$. Therefore:

$$\frac{Q(z_i^0; c, w)}{Q(z_i^1; c, w)} = \delta_i \cdot \left(\frac{x_i^0}{x_i^1}\right) \tag{38}$$

which relates the relative transmission efficiency of adjacent precursor isotopes to the distortion between observed and theoretical isotope ratios.

After assigning a functional form to $Q$ it is possible to estimate the parameters of the quadrupole transmission function given the observations. We propose a solution in Section 1.14.2. Section 1.14.1 describes how to estimate each $x_i^k$.

##### 1.14.1 Estimating Observed Precursor Isotope Abundances

Let $p(k)$ denote the $k$-th isotope of a precursor ion species, $p$, and then let $s\left(f(l)_j^{p(k)}\right)$ give the conditional abundance of $l$-th isotope of the $j$-th fragment of $p(k)$. Then construct the following matrix:

$$\mathbf{S}_p = \begin{bmatrix} s\left(f(0)_1^{p(0)}\right) & s\left(f(0)_1^{p(1)}\right) & \dots & s\left(f(0)_1^{p(n)}\right) \\ s\left(f(1)_1^{p(0)}\right) & s\left(f(1)_1^{p(1)}\right) & \dots & s\left(f(1)_1^{p(n)}\right) \\ s\left(f(2)_1^{p(0)}\right) & s\left(f(2)_1^{p(1)}\right) & \dots & s\left(f(2)_1^{p(n)}\right) \\ s\left(f(0)_2^{p(0)}\right) & s\left(f(0)_2^{p(1)}\right) & \dots & s\left(f(0)_2^{p(n)}\right) \\ \vdots & \vdots & \ddots & \vdots \\ s\left(f(2)_m^{p(0)}\right) & s\left(f(2)_m^{p(1)}\right) & \dots & s\left(f(2)_m^{p(n)}\right) \end{bmatrix} \tag{39}$$

Now for an observed MS2 spectrum, let $\vec{I}_i$ be the observed intensities of peaks that match each fragment isotope, with a value of zero for unmatched fragment ions. This results in the following linear system:

$$\mathbf{S}_p \begin{bmatrix} y^0 \\ y^1 \\ \vdots \\ y^n \end{bmatrix} \approx \vec{I}_i \tag{40}$$

The least squares solution, $\vec{y}$, approximates the relative abundances of the isotopes of the precursor $p$ after quadrupole transmission.

##### 1.14.2 Estimating Quadrupole Transmission with an Asymmetric Generalized Bell Function

Pioneer models quadrupole transmission efficiency using an asymmetric generalized bell function, a modified form of the generalized bell function [15].

$$Q(x; a_l, a_r, b_l, b_r) = \begin{cases} \left[1 + (-x \,/\, a_l)^{2b_l}\right]^{-1} & \text{if } x \leq 0 \\ \left[1 + (x \,/\, a_r)^{2b_r}\right]^{-1} & \text{if } x > 0 \end{cases} \tag{41}$$

In this four-parameter model, $a_l$ and $a_r$ control the isolation window width. They represent the distance from the maximum peak (at $x = 0$) to the half-maximum intensity on the left and right sides, respectively. The shape parameters $b_l$ and $b_r$ determine the smoothness of the transition. Given that $x^0 < x^1$, the ratio of the transmission function at $x^0$ and $x^1$ is as follows.

$$R(x_i^0, x_i^1) = \frac{Q(x_i^0; a_l, a_r, b_l, b_r)}{Q(x_i^1; a_l, a_r, b_l, b_r)}. \tag{42}$$

Expanding piecewise:

$$R(x_i^0, x_i^1) = \begin{cases} \left(\frac{1+\left(-x^1 \,/\, a_l\right)^{2b_l}}{1+(-x^0 \,/\, a_l)^{2b_l}}\right) & \text{if } x^0 \leq 0 \text{ and } x^1 \leq 0 \\ \left(\frac{1+\left(z^1 \,/\, a_l\right)^{2b_l}}{1+(-x^0 \,/\, a_r)^{2b_r}}\right) & \text{if } x^0 \leq 0 \text{ and } x^1 > 0 \\ \left(\frac{1+\left(x^1 \,/\, a_r\right)^{2b_r}}{1+(x^0 \,/\, a_r)^{2b_r}}\right) & \text{if } x^0 > 0 \text{ and } x^1 > 0 \end{cases} \tag{43}$$

Therefore, to estimate the parameters, $a_l, a_r, b_l, b_r$, Pioneer minimizes the squared error between the respective natural logarithms of $R(x_0, x_1)$ and the data as in equation (1.14). The objective function to minimize becomes:

$$\min_{a_l, a_r, b_l, b_r} L(a_l, a_r, b_l, b_r; \vec{x^0}, \vec{x^1}) = \sum_i \left( \ln\left(R(x_i^0, x_i^1)\right) - \ln\left(\delta_i \cdot \frac{x_i^0}{x_i^1}\right)\right)^2 \tag{44}$$

Pioneer uses the Symbolics.jl package in Julia to calculate the piecewise partial derivatives for $R$ with respect to the parameters and then implements a Levenberg-Marquardt algorithm to minimize the objective function, $L$.

### 2 Supplementary Figures

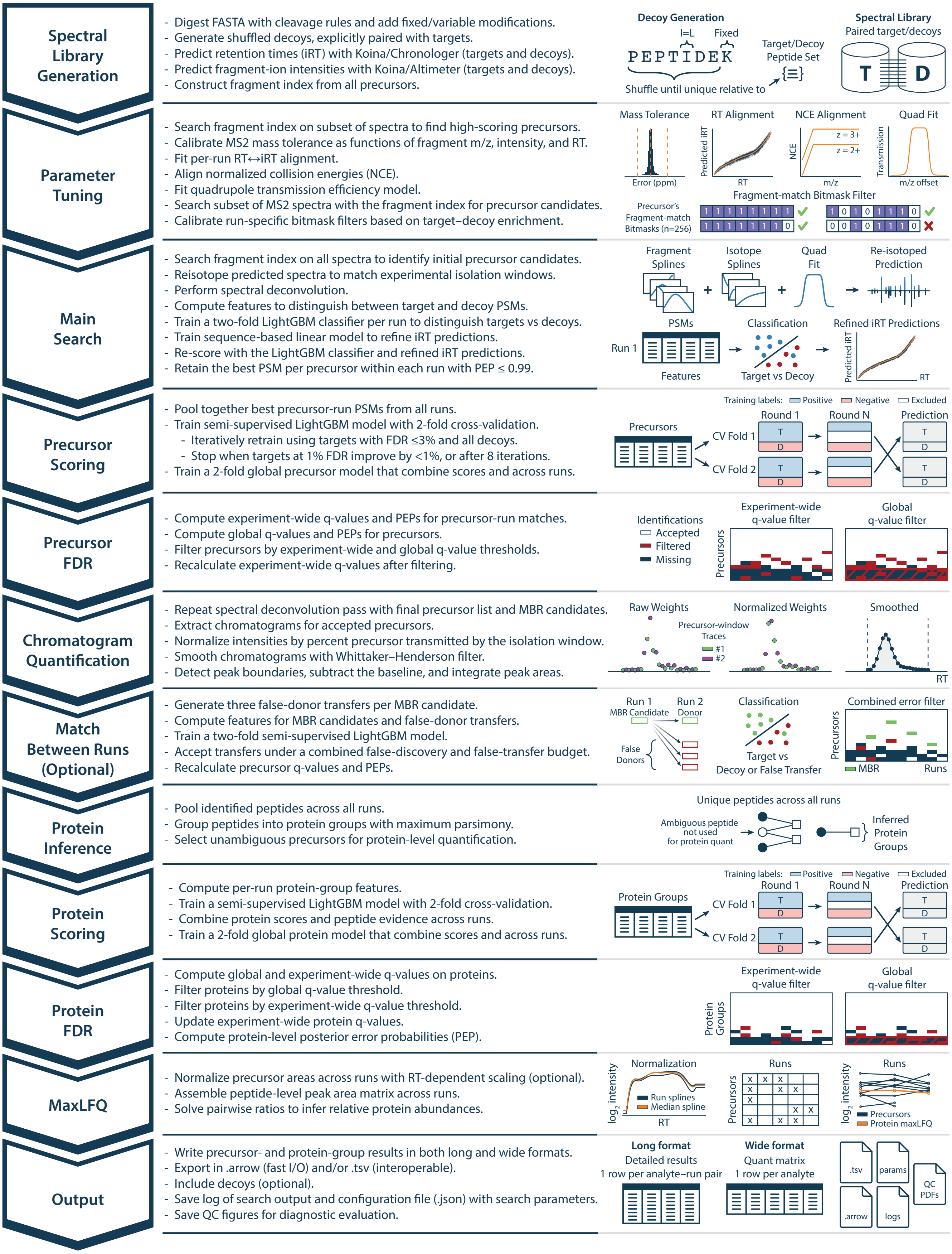


**Supplementary Fig. 1. Detailed Pioneer workflow for DIA analysis.** Each module of the pipeline is shown with its major computational steps (left) and schematic illustrations (right).

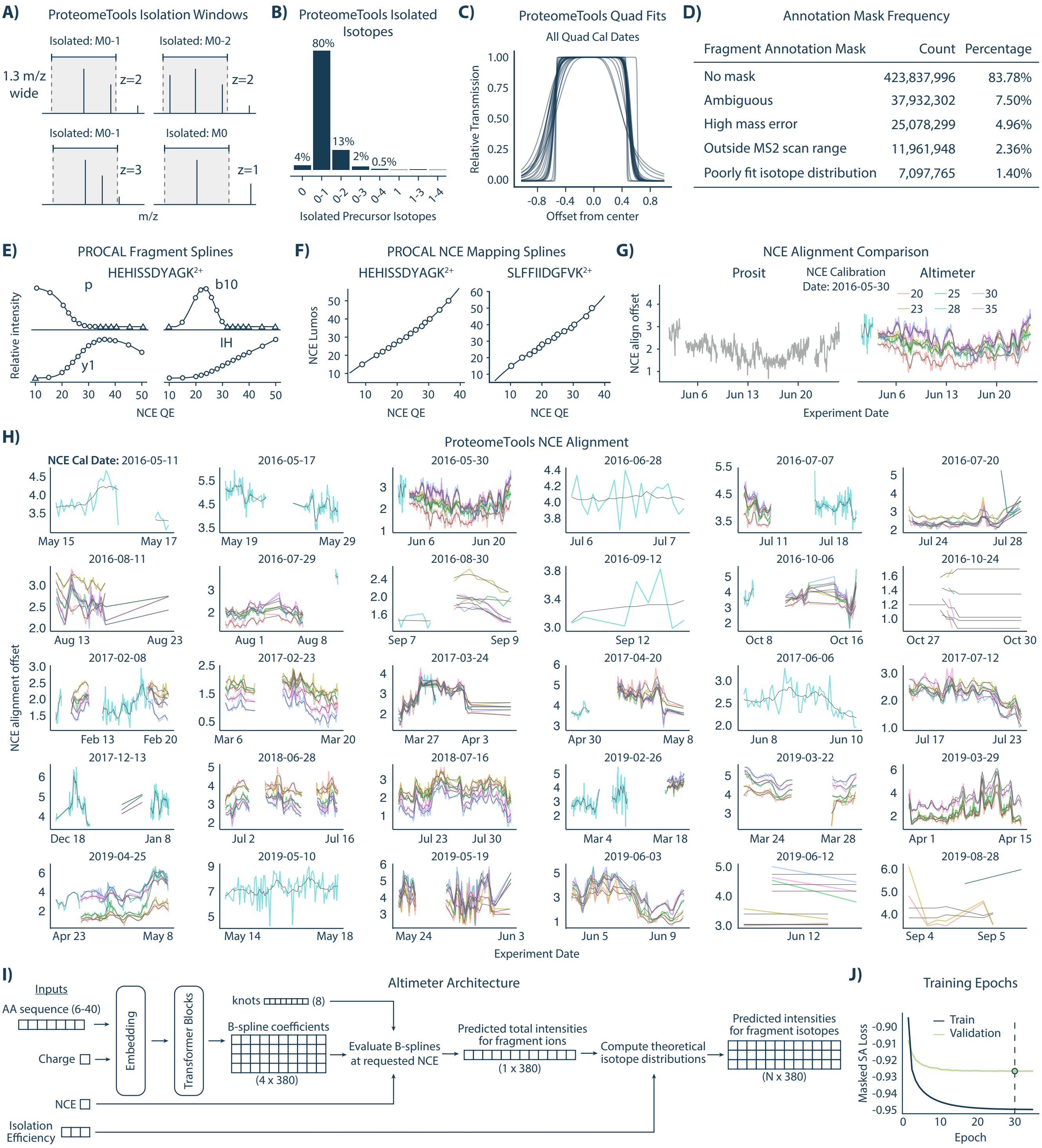


**Extended Data Fig. 2. Training data calibration, dataset characterization, and training details for Altimeter.**
**A)** Examples of isolation windows in the ProteomeTools dataset, illustrating how isolated isotopes depend on window width and precursor charge.
**B)** Distribution of precursor isotopes isolated in ProteomeTools assuming a perfect box shape for transmission.
**C)** Fitted quadrupole tranmission profiles across all ProteomeTools quadrupole calibration dates.
**D)** Frequency of fragment annotation mask types across the training dataset.
**E)** Example B-spline fits to PROCAL peptide fragment intensities across 15 normalized collision energies (NCEs) acquired on a Q Exactive Plus.
**F)** PROCAL peptide spectra aligned between a Lumos and QE Plus, with calibration curves generated for individual precursors.
**G)** Smoothed NCE alignment offsets (dark gray) for different NCE settings compared to Prosit. Alignment offset is relative to the NCE used for acquisition.
**H)** Smoothed NCE alignment offsets (dark gray) across the ProteomeTools dataset over runs, grouped by NCE calibration dates.
**I)** Altimeter architecture: precursor sequence and charge are input to a transformer, which outputs four B-spline coefficients per fragment. Shared knots are evaluated at given NCEs to generate predicted spectra, where losses are computed. Isolation efficiency can be included to model isotope distributions.
**J)** Training and validation loss across epochs, showing model convergence. The model with minimal validation loss was chosen (dashed line).

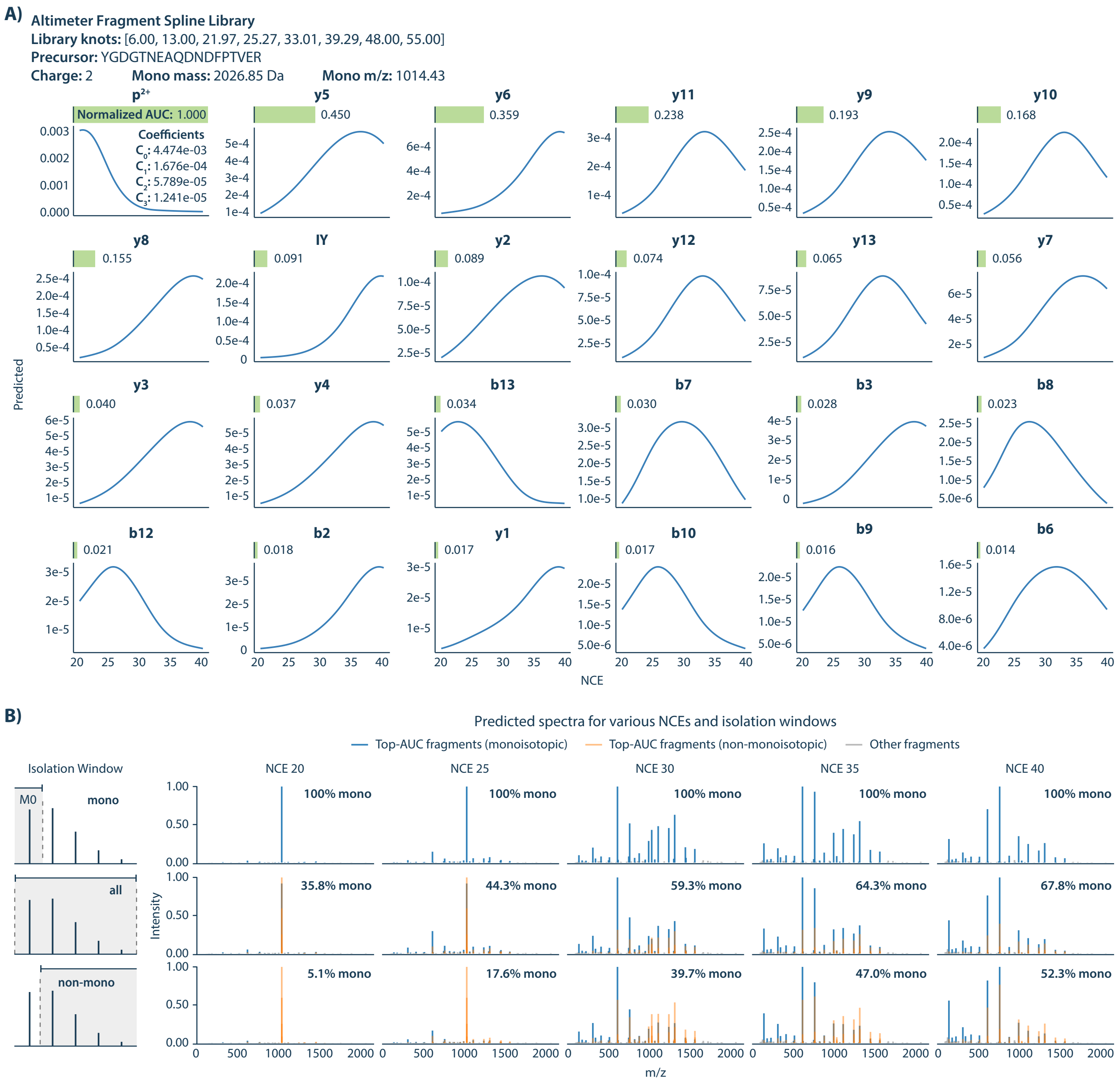


**Extended Data Fig. 3. Fragment splines and predicted spectra from Altimeter.**
**A)** Example fragment spline library for the peptide YGDGTNEAQDNDFPTVER$^{2+}$. Each fragment ion is modeled with four B-spline coefficients (inset, $p^{2+}$), evaluated across normalized collision energies (NCEs). Area under the curve (AUC) values are normalized by the most abundant AUC and provide a measure of fragment abundance across NCEs. The 24 highest-AUC fragments are shown. Pioneer does not use precursor or immonium ions unless requested.
**B)** Predicted spectra for the same peptide at NCE values from 20–40, under different isolation window assumptions: monoisotopic only (top), all isotopes (middle), and non-monoisotopic only (bottom). All fragments are displayed, with high-AUC fragments from panel A highlighted (blue: monoisotopes; orange: non-monoisotopes). Percentages indicate the fraction of predicted MS2 intensity arising from monoisotopic fragments.

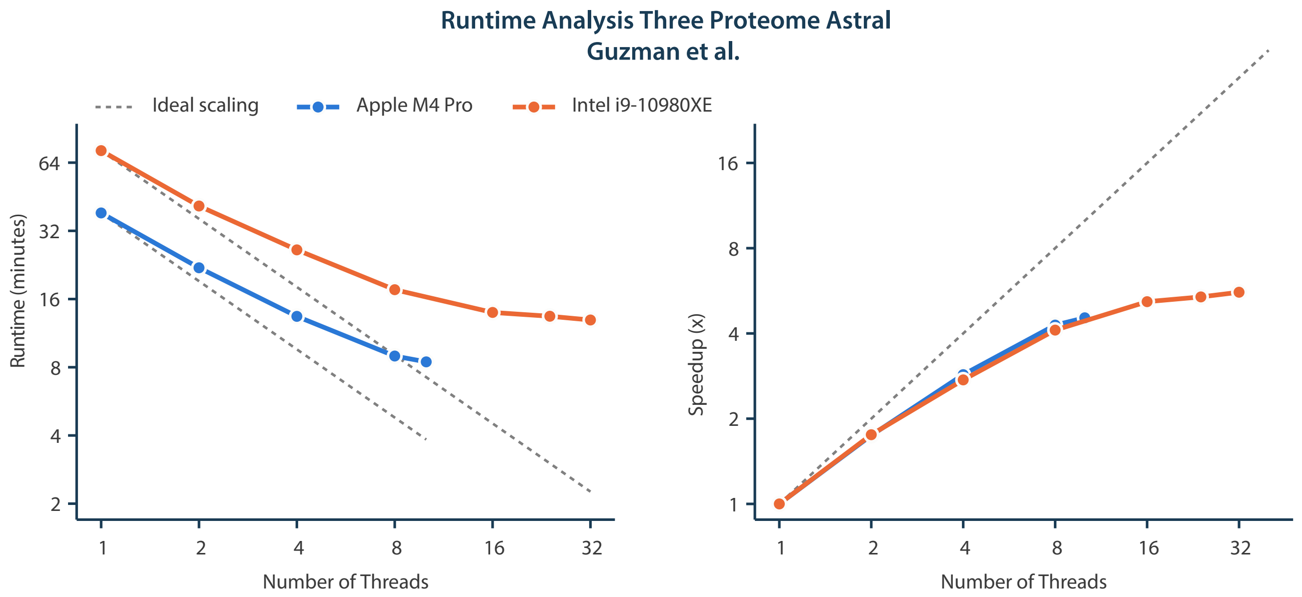


**Extended Data Fig. 5. Runtime Scaling with Respect to the Number of Threads**

**A)** Pioneer runtime as a function of CPU thread count for the three-proteome Astral dataset from Guzman et al. Left, total runtime; right, speedup relative to the single-thread case. The dashed gray line indicates ideal linear scaling. The blue curves represent an analysis performed on an Apple MacBook Pro with an M4 Pro chip (10 performance + 4 efficiency cores) and 48 GB RAM. The orange curves represent an analysis performed on a Dell Precision 5820 Tower workstation with an Intel Core i9-10980XE processor (18 physical cores, 36 threads, 3.00 GHz base) and 64 GB RAM, running Windows 11.

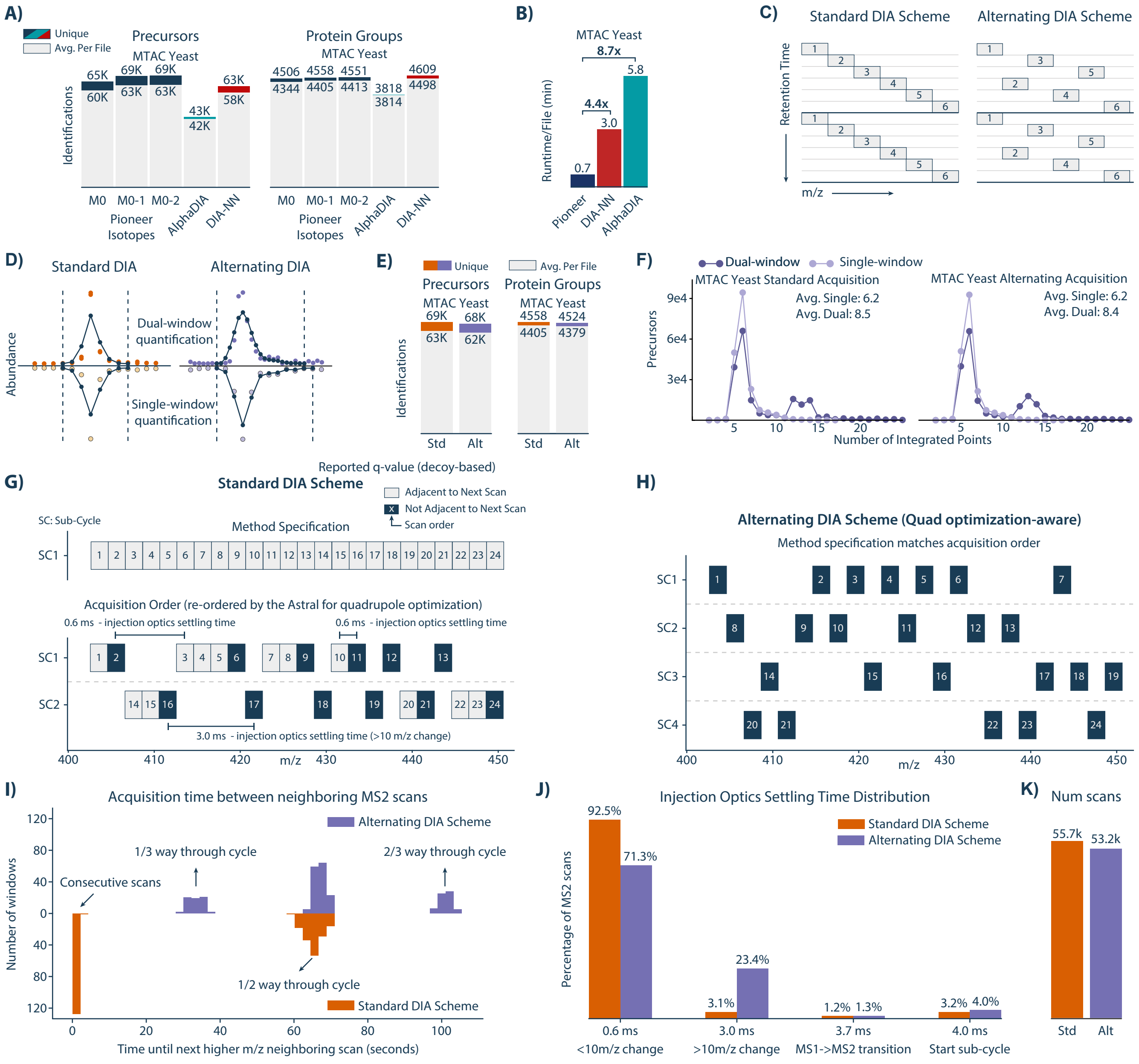


**Extended Data Fig. 4. DIA Acquisition Methods.**

**A)** Precursor and protein group identifications from the MTAC yeast experiment using different numbers of fragment isotopes for Pioneer and compared to AlphaDIA and DIA-NN.

**B)** Search Runtimes from the MTAC yeast experiment for Pioneer, DIA-NN, and AlphaDIA.

**C)** Schematic of standard and alternating DIA acquisition schemes.

**D)** Example chromatograms for the same precursor across the combinations of acquisition (standard/alternating) and quantification (daul-/single-window) methods.

**E)** Precursor and protein group identifications from the MTAC yeast experiment for the standard (Std) and alternating (Alt) window acquisition methods.

**F)** Distribution of integrated points per precursor for dual- versus single-window quantification.

**G-H)** Internal optimizations performed by the instrument control software rearrange the specified acquisition window order. Panel G shows the order in which the windows are specified for a representitive portion of the cycle (top). The bottom panel shows the order in which the scans are actually performed. Panel H shows the order in which the scans were performed for the alternating window method.

**I)** Histogram of the times in seconds between neighboring MS2 scans. Neighboring scans are defined as those with adjacent m/z values. The top (purple) is the histogram for the alternating window method and the bottom (orange) is the histogram for the standard window method.

**J)** Bar plot of injection optics settling times for the alternating and standard window methods. The x-axis indicates the time and the condition under which that time occurs. The y-axis represents the percentage of total scans falling into each x-axis bin.

**K)** Bar plot of the total number of scans per run for the standard and alternating window methods.

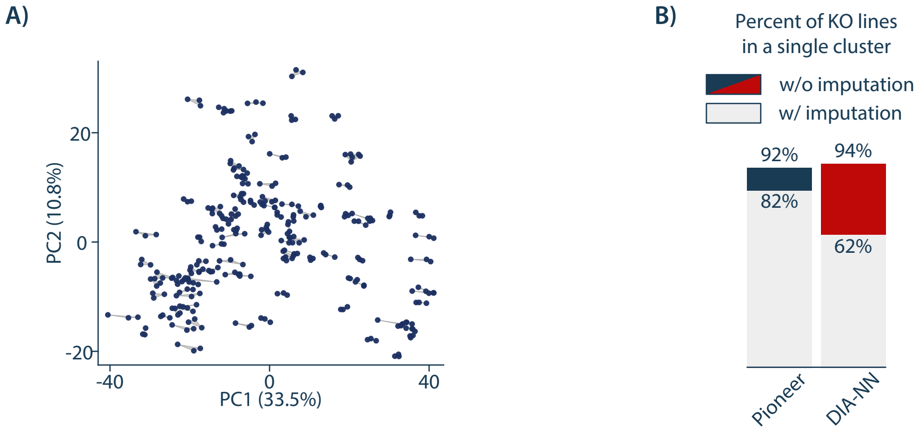


**Extended Data Fig. 6. Replicate Consistency**
**A)** PCA analysis of the Pioneer yeast KO runs after imputation. Imputation was performed in Perseus with default parameters (width 0.3, downshift 1.8) after deleting rows that were not complete for at least one knockout or control condition. **B)** Agglomerative clustering of the Pioneer yeast KO runs was performed after imputation. Clustering was performed using the ward method and for a predetermined number of clusters equal to the number (104) of knockout conditions and controls. Bar heights show the percentages of conditions where all three replicates were found in the same cluster when clustering was performed before (colored) or after (grey) imputation.

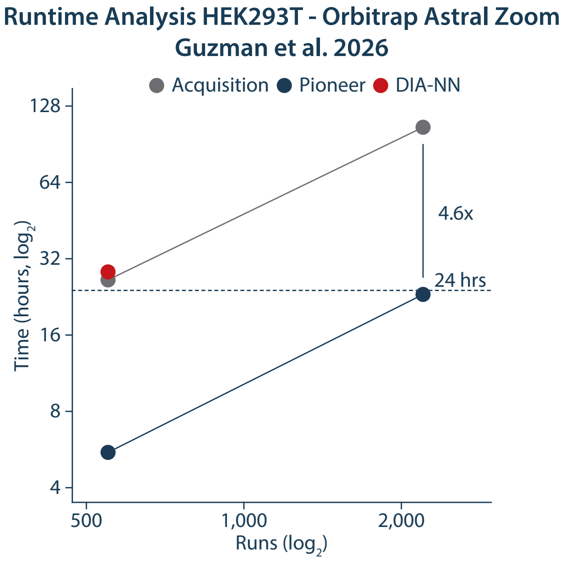


**Extended Data Fig. 7. Runtime Scaling with Respect to the Number of Runs**

Pioneer v0.12.0 and DIA-NN v2.6.0 runtime as a function of the number of runs, using the 500 samples-per-day (SPD) HEK293T dataset acquired on an Orbitrap Astral Zoom from Guzman et al. 2026 (PXD071972; 550 runs). The number of runs was increased to 2,200 using symbolic links to the original files. Navy points show Pioneer runtimes, and the red point shows DIA-NN runtime. Grey points show the instrument acquisition time for the same number of runs. Each search had a dedicated node with two Intel Xeon Gold 6548Y+ processors (32 cores each, 2.5 GHz base). Each search was pinned to 16-cores. Pioneer was limited to 64 GB of RAM. DIA-NN ran out of memory at 64 GB for 550 runs, so DIA-NN's allotment was increased to 128 GB.

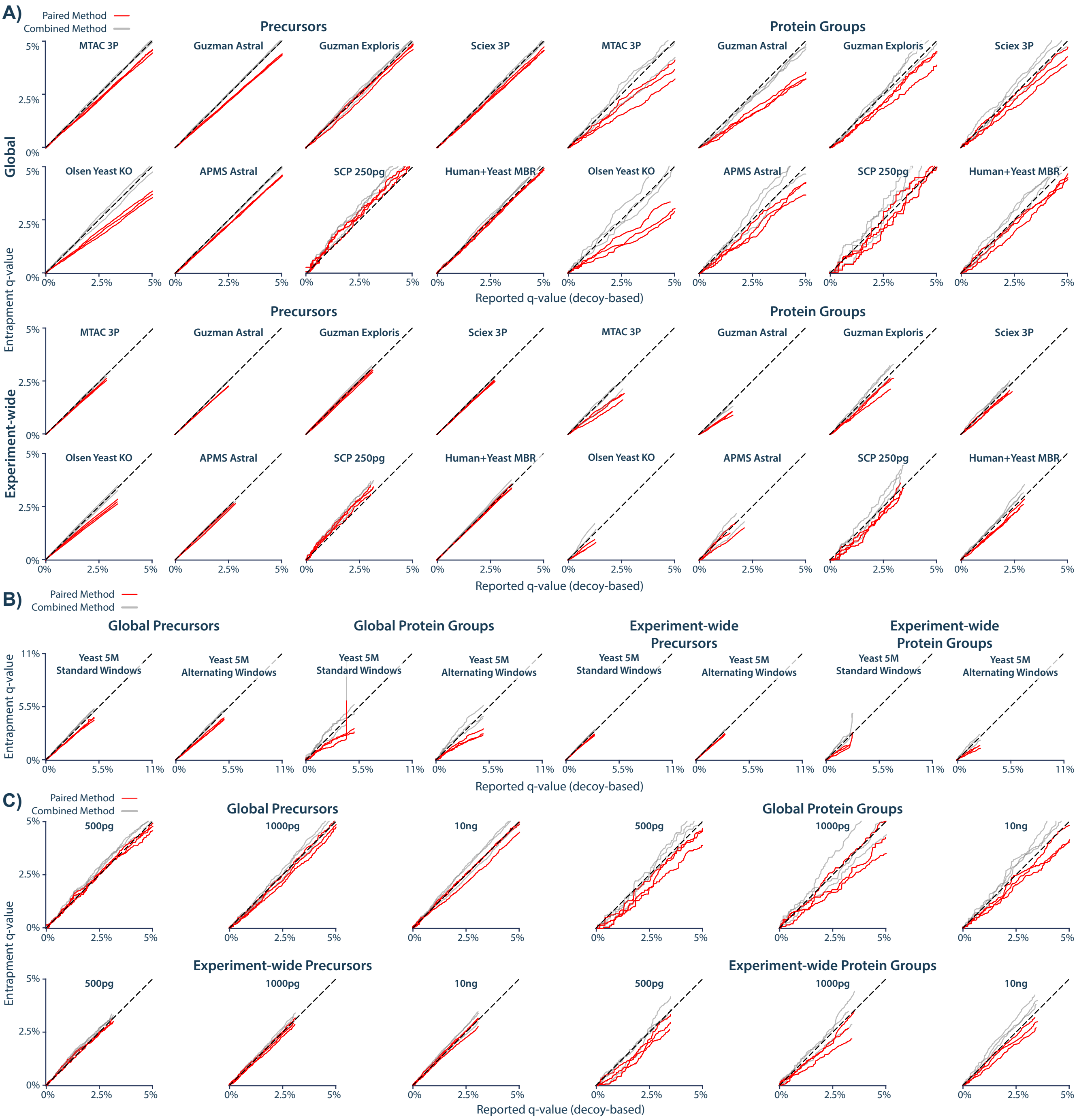


**Extended Data Fig. 8. Entrapment analysis at relaxed stringency.**

**A)** Entrapment analyses for Pioneer searches repeated from Figure 5C of the main manuscript with a q-value threshold of 5%. Analyses were conducted three times with three independently generated entrapment libraries.

**B)** Triplicate entrapment analyses for Pioneer searches of the MTAC Yeast experiment (Figure 3A) searches with a q-value threshold of 5%. This includes the standard and alternating window acquisition versions.

**C)** Triplicate entrapment analysis for the single cell equivalent experiment from Petrosius et al. 2025 which are also shown in Figure 4. Searches were repeated with a q-value threshold of 5%.
