## Extended Data Figure 1 for "Pioneer and Altimeter: Rapid and Scalable Analysis of DIA Proteomics Data Optimized for Narrow Isolation Windows"

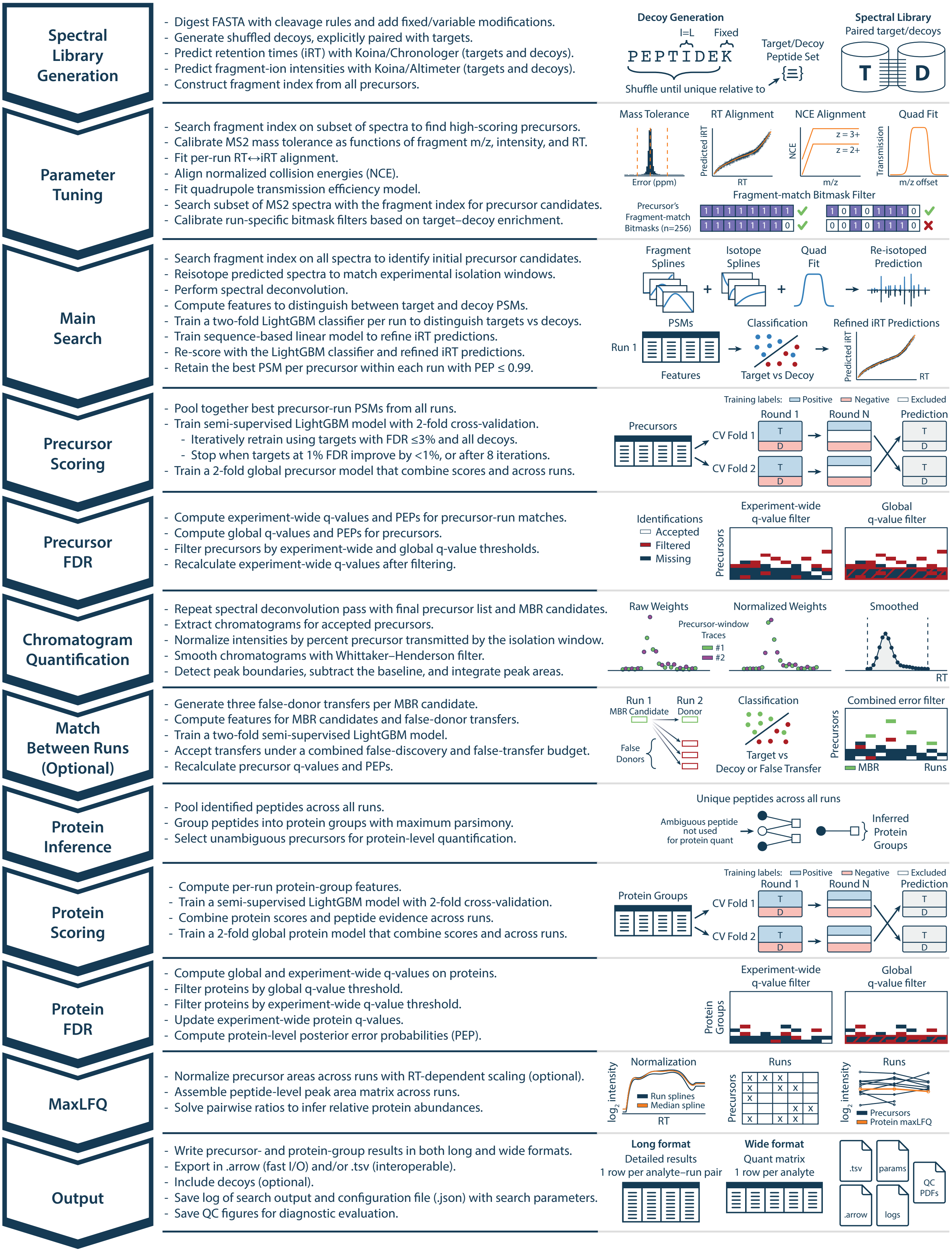


**Supplementary Fig. 1. Detailed Pioneer workflow for DIA analysis.** Each module of the pipeline is shown with its major computational steps (left) and schematic illustrations (right).
