## Extended Data Figure 2 for "Pioneer and Altimeter: Rapid and Scalable Analysis of DIA Proteomics Data Optimized for Narrow Isolation Windows"

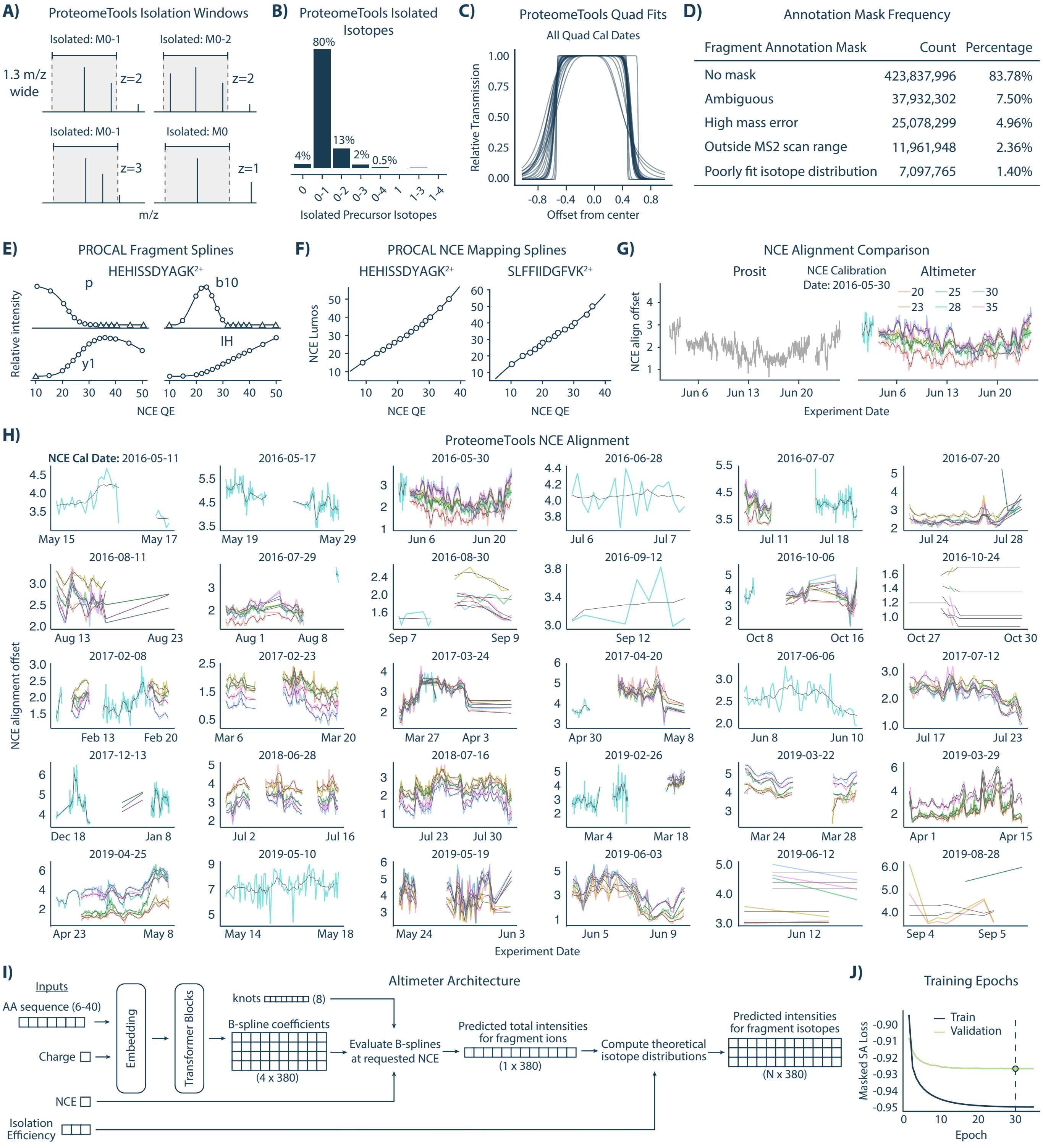


**Extended Data Fig. 2. Training data calibration, dataset characterization, and training details for Altimeter.**
**A)** Examples of isolation windows in the ProteomeTools dataset, illustrating how isolated isotopes depend on window width and precursor charge.
**B)** Distribution of precursor isotopes isolated in ProteomeTools assuming a perfect box shape for transmission.
**C)** Fitted quadrupole tranmission profiles across all ProteomeTools quadrupole calibration dates.
**D)** Frequency of fragment annotation mask types across the training dataset.
**E)** Example B-spline fits to PROCAL peptide fragment intensities across 15 normalized collision energies (NCEs) acquired on a Q Exactive Plus.
**F)** PROCAL peptide spectra aligned between a Lumos and QE Plus, with calibration curves generated for individual precursors.
**G)** Smoothed NCE alignment offsets (dark gray) for different NCE settings compared to Prosit. Alignment offset is relative to the NCE used for acquisition.
**H)** Smoothed NCE alignment offsets (dark gray) across the ProteomeTools dataset over runs, grouped by NCE calibration dates.
**I)** Altimeter architecture: precursor sequence and charge are input to a transformer, which outputs four B-spline coefficients per fragment. Shared knots are evaluated at given NCEs to generate predicted spectra, where losses are computed. Isolation efficiency can be included to model isotope distributions.
**J)** Training and validation loss across epochs, showing model convergence. The model with minimal validation loss was chosen (dashed line).
