## Extended Data Figure 3 for "Pioneer and Altimeter: Rapid and Scalable Analysis of DIA Proteomics Data Optimized for Narrow Isolation Windows"

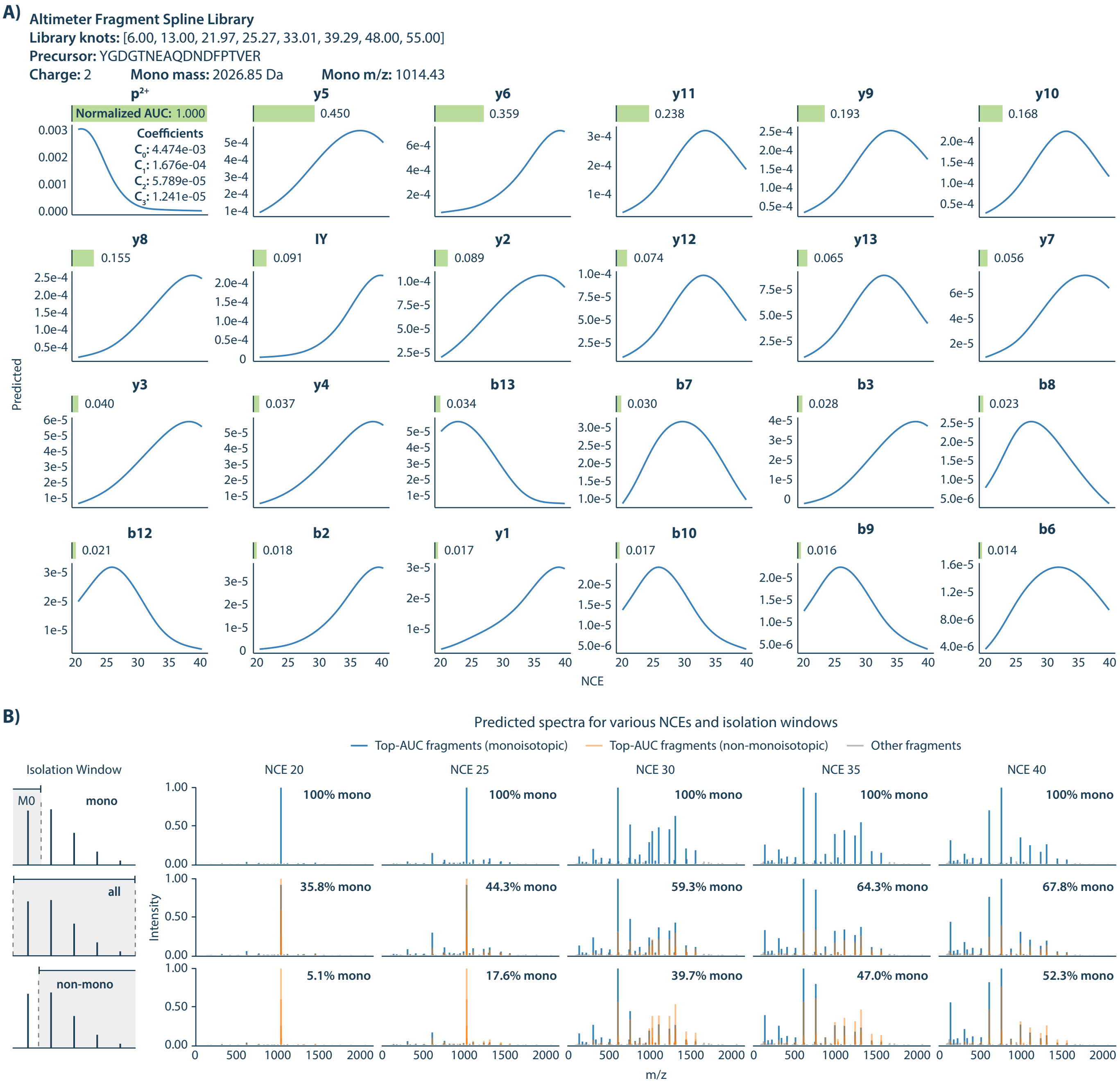


**Extended Data Fig. 3. Fragment splines and predicted spectra from Altimeter.**

**A)** Example fragment spline library for the peptide YGDGTNEAQDNDFPTVER$^{2+}$. Each fragment ion is modeled with four B-spline coefficients (inset, $p^{2+}$), evaluated across normalized collision energies (NCEs). Area under the curve (AUC) values are normalized by the most abundant AUC and provide a measure of fragment abundance across NCEs. The 24 highest-AUC fragments are shown. Pioneer does not use precursor or immonium ions unless requested.

**B)** Predicted spectra for the same peptide at NCE values from 20–40, under different isolation window assumptions: monoisotopic only (top), all isotopes (middle), and non-monoisotopic only (bottom). All fragments are displayed, with high-AUC fragments from panel A highlighted (blue: monoisotopes; orange: non-monoisotopes). Percentages indicate the fraction of predicted MS2 intensity arising from monoisotopic fragments.
