## Extended Data Figure 4 for "Pioneer and Altimeter: Rapid and Scalable Analysis of DIA Proteomics Data Optimized for Narrow Isolation Windows"

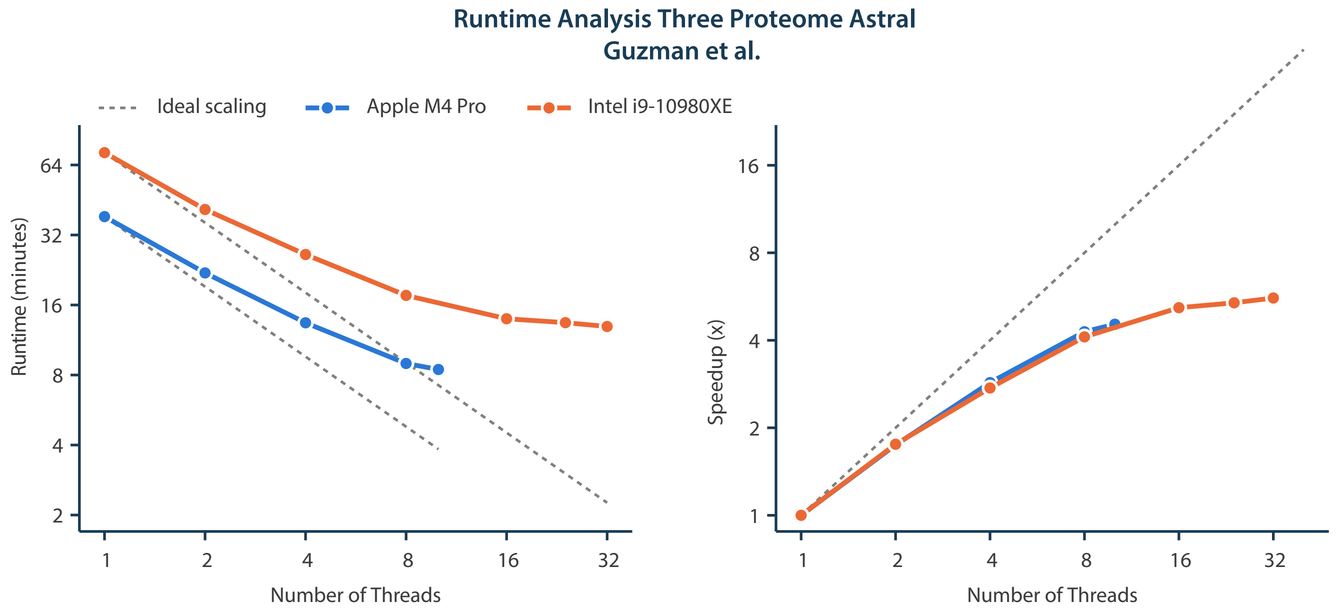


**Extended Data Fig. 5. Runtime Scaling with Respect to the Number of Threads**
**A)** Pioneer runtime as a function of CPU thread count for the three-proteome Astral dataset from Guzman et al. Left, total runtime; right, speedup relative to the single-thread case. The dashed gray line indicates ideal linear scaling. The blue curves represent an analysis performed on an Apple MacBook Pro with an M4 Pro chip (10 performance + 4 efficiency cores) and 48 GB RAM. The orange curves represent an analysis performed on a Dell Precision 5820 Tower workstation with an Intel Core i9-10980XE processor (18 physical cores, 36 threads, 3.00 GHz base) and 64 GB RAM, running Windows 11.
