## Extended Data Figure 5 for "Pioneer and Altimeter: Rapid and Scalable Analysis of DIA Proteomics Data Optimized for Narrow Isolation Windows"

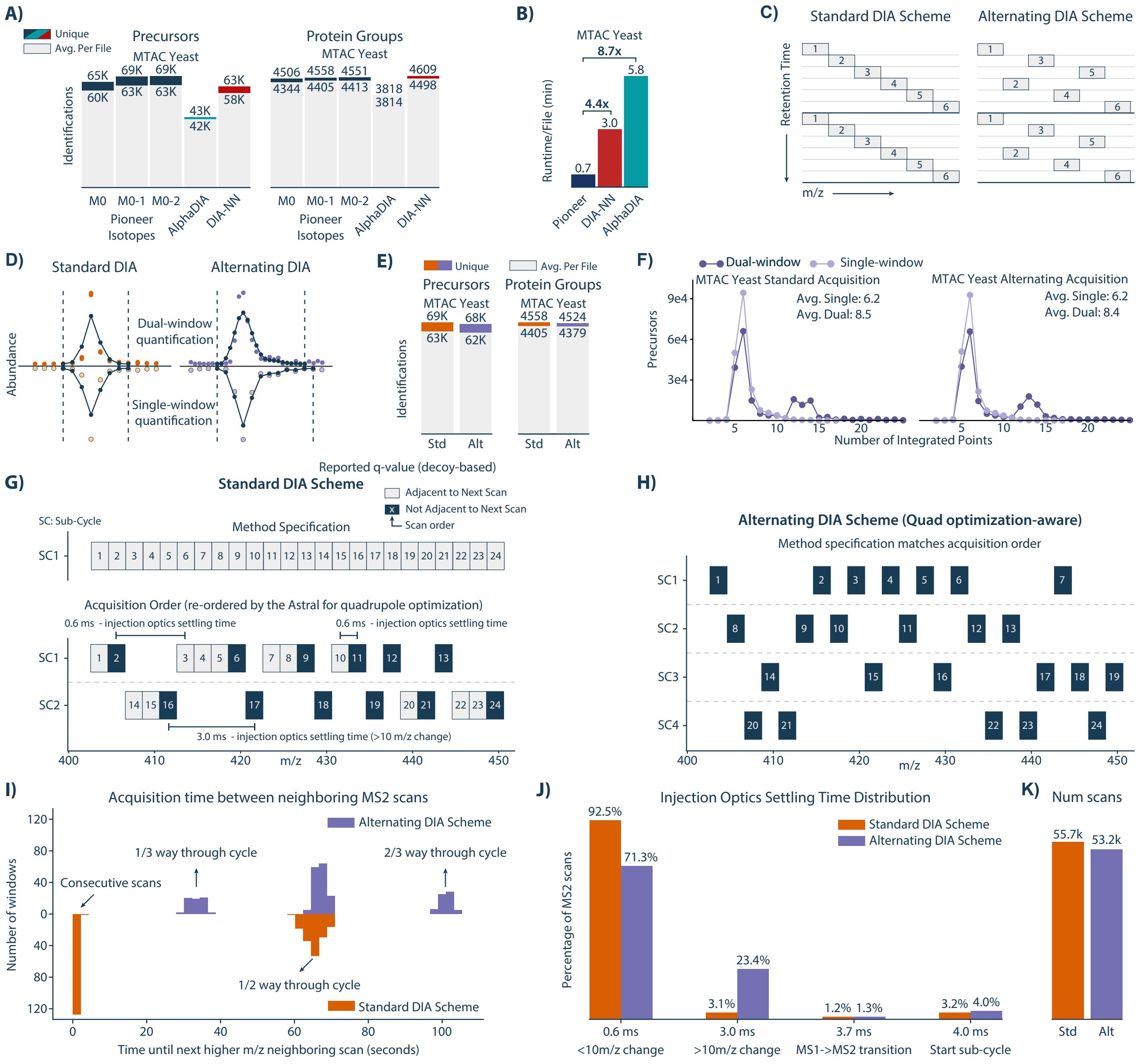


**Extended Data Fig. 4. DIA Acquisition Methods.**

**A)** Precursor and protein group identifications from the MTAC yeast experiment using different numbers of fragment isotopes for Pioneer and compared to AlphaDIA and DIA-NN.

**B)** Search Runtimes from the MTAC yeast experiment for Pioneer, DIA-NN, and AlphaDIA.

**C)** Schematic of standard and alternating DIA acquisition schemes.

**D)** Example chromatograms for the same precursor across the combinations of acquisition (standard/alternating) and quantification (daul-/single-window) methods.

**E)** Precursor and protein group identifications from the MTAC yeast experiment for the standard (Std) and alternating (Alt) window acquisition methods.

**F)** Distribution of integrated points per precursor for dual- versus single-window quantification.

**G-H)** Internal optimizations performed by the instrument control software rearrange the specified acquisition window order. Panel G shows the order in which the windows are specified for a representitive portion of the cycle (top). The bottom panel shows the order in which the scans are actually performed. Panel H shows the order in which the scans were performed for the alternating window method.

**I)** Histogram of the times in seconds between neighboring MS2 scans. Neighboring scans are defined as those with adjacent m/z values. The top (purple) is the histogram for the alternating window method and the bottom (orange) is the histogram for the standard window method.

**J)** Bar plot of injection optics settling times for the alternating and standard window methods. The x-axis indicates the time and the condition under which that time occurs. The y-axis represents the percentage of total scans falling into each x-axis bin.

**K)** Bar plot of the total number of scans per run for the standard and alternating window methods.
