## Extended Data Figure 6 for "Pioneer and Altimeter: Rapid and Scalable Analysis of DIA Proteomics Data Optimized for Narrow Isolation Windows"

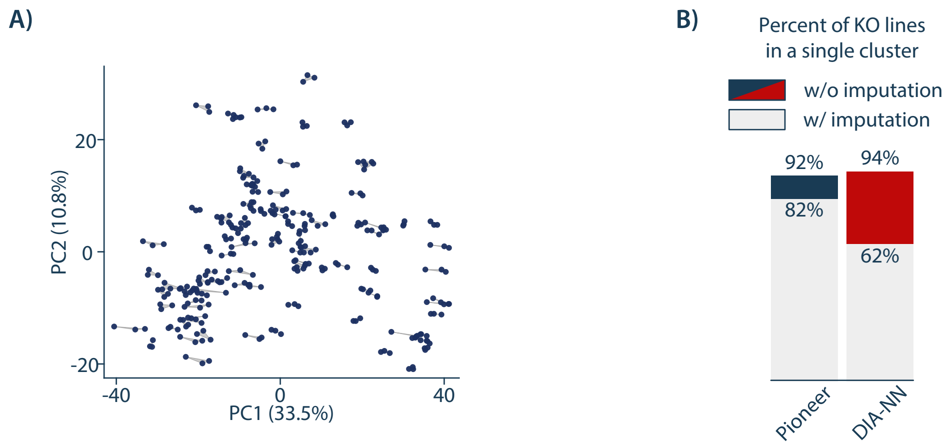


**Extended Data Fig. 6. Replicate Consistency**

**A)** PCA analysis of the Pioneer yeast KO runs after imputation. Imputation was performed in Perseus with default parameters (width 0.3, downshift 1.8) after deleting rows that were not complete for at least one knockout or control condition. **B)** Agglomerative clustering of the Pioneer yeast KO runs was performed after imputation. Clustering was performed using the ward method and for a predetermined number of clusters equal to the number (104) of knockout conditions and controls. Bar heights show the percentages of conditions where all three replicates were found in the same cluster when clustering was performed before (colored) or after (grey) imputation.
