## Extended Data Figure 7 for "Pioneer and Altimeter: Rapid and Scalable Analysis of DIA Proteomics Data Optimized for Narrow Isolation Windows"

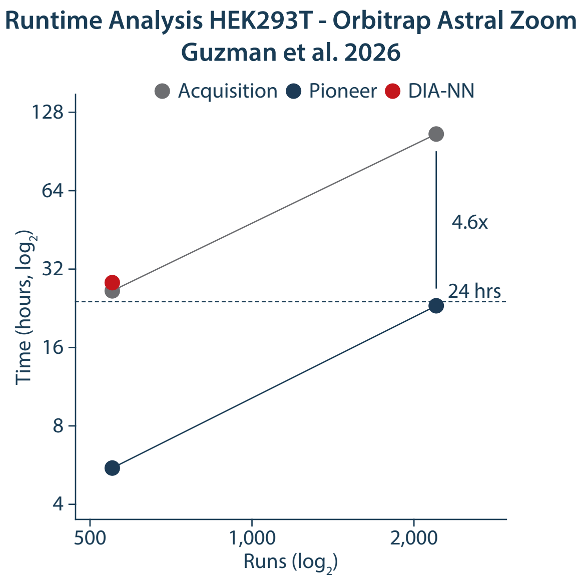


**Extended Data Fig. 7. Runtime Scaling with Respect to the Number of Runs**

Pioneer v0.12.0 and DIA-NN v2.6.0 runtime as a function of the number of runs, using the 500 samples-per-day (SPD) HEK293T dataset acquired on an Orbitrap Astral Zoom from Guzman et al. 2026 (PXD071972; 550 runs). The number of runs was increased to 2,200 using symbolic links to the original files. Navy points show Pioneer runtimes, and the red point shows DIA-NN runtime. Grey points show the instrument acquisition time for the same number of runs. Each search had a dedicated node with two Intel Xeon Gold 6548Y+ processors (32 cores each, 2.5 GHz base). Each search was pinned to 16-cores. Pioneer was limited to 64 GB of RAM. DIA-NN ran out of memory at 64 GB for 550 runs, so DIA-NN's allotment was increased to 128 GB.
