## Extended Data Figure 8 for "Pioneer and Altimeter: Rapid and Scalable Analysis of DIA Proteomics Data Optimized for Narrow Isolation Windows"

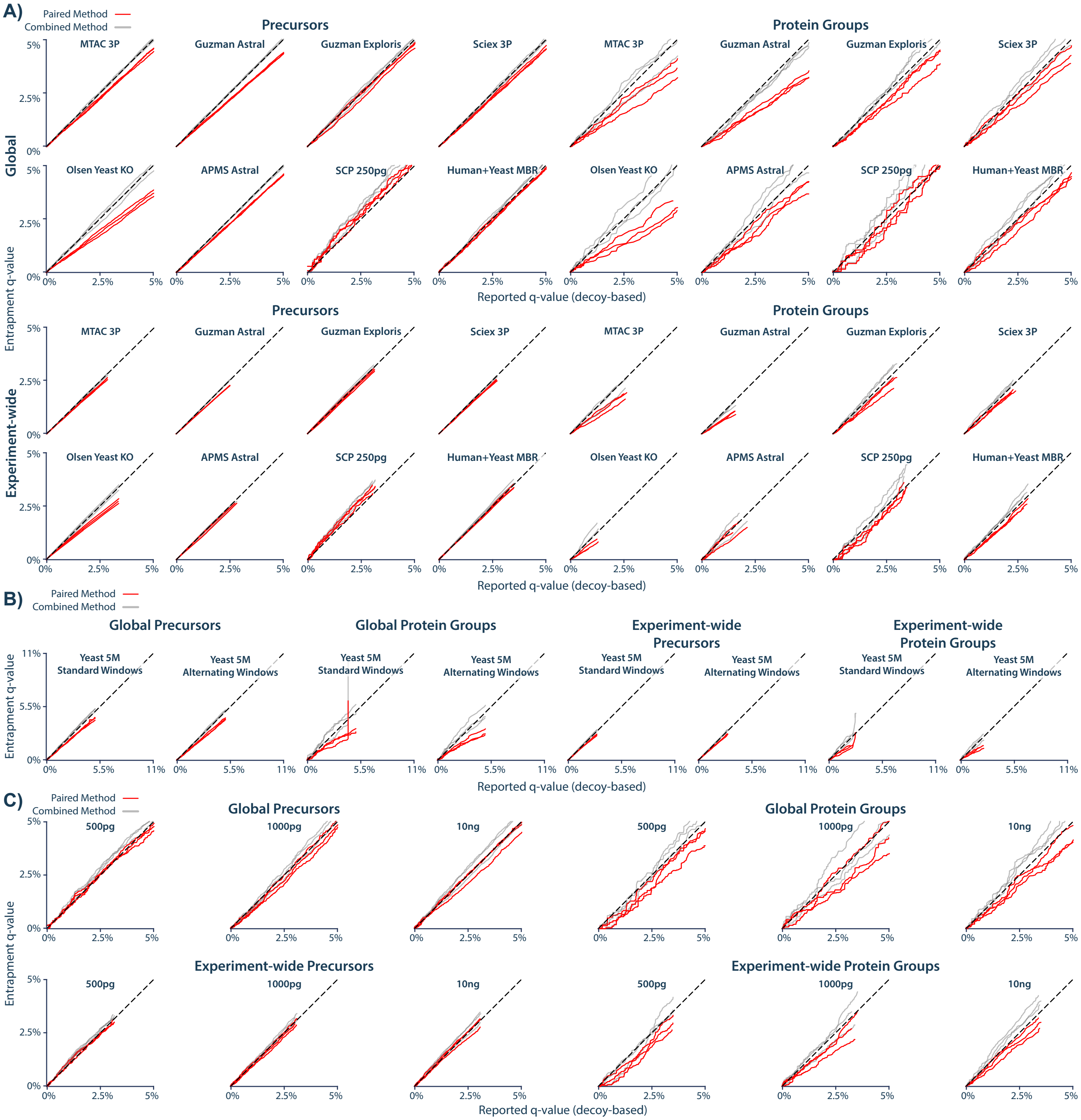


**Extended Data Fig. 8. Entrapment analysis at relaxed stringency.**

**A)** Entrapment analyses for Pioneer searches repeated from Figure 5C of the main manuscript with a q-value threshold of 5%. Analyses were conducted three times with three independently generated entrapment libraries.

**B)** Triplicate entrapment analyses for Pioneer searches of the MTAC Yeast experiment (Figure 3A) searches with a q-value threshold of 5%. This includes the standard and alternating window acquisition versions.

**C)** Triplicate entrapment analysis for the single cell equivalent experiment from Petrosius et al. 2025 which are also shown in Figure 4. Searches were repeated with a q-value threshold of 5%.
